# Mechanistic divergence of piRNA biogenesis in Drosophila

**DOI:** 10.1101/2022.11.14.516378

**Authors:** Shashank Chary, Rippei Hayashi

## Abstract

Organisms require mechanisms to distinguish self and non-self RNA. This distinction is crucial to initiate the biogenesis of piRNAs. In Drosophila ovaries, PIWI-guided slicing and the recognition of piRNA precursor transcripts by the DEAD-box RNA helicase Yb are the two known mechanisms to licence an RNA for piRNA biogenesis in the germline and the soma, respectively. Both, the PIWI proteins and Yb are highly conserved across most Drosophila species and are thought to be essential to the piRNA pathway and for silencing transposons. However, we find that species closely related to *D. melanogaster* have lost the *yb* gene, as well as the PIWI gene *Ago3*. We show that the precursor RNA is still selected in the absence of Yb to abundantly generate transposon antisense piRNAs in the soma. We further demonstrate that *D. eugracilis*, which lacks *Ago3*, is completely devoid of ping-pong piRNAs and exclusively produces phased piRNAs in the absence of slicing. Thus, there are more possible routes through which the piRNA pathway can achieve specificity than previously suggested.

## Introduction

The everchanging landscape of transposable elements (TEs) in the genome requires the host defence pathway to be highly plastic and adaptive to maintain silencing. One such mechanism is the Piwi-interacting RNA (piRNA) silencing pathway. piRNA is a class of small RNA of 25 to 32 nucleotides in length and is loaded onto the PIWI-clade Argonaute proteins to form the piRNA-induced silencing complex (pi-RISC)^1, 2^. piRNAs are highly diverse in sequence yet are highly specific to TE sequences. Mechanisms underlying the distinction between TEs and host genes remain to be fully understood.

The piRNA pathway is predominantly active in the germline lineage, protecting genome integrity and gonadal development from harmful TE insertions^3^. Some TEs can be uniquely expressed in somatic cells and can carry genes that allow them to infect other cells^4, 5^. One notable example are *gypsy* retrotransposons in Drosophila that carry retroviral envelope (*env*) genes^6^, which are expressed in the ovarian somatic cells and can infect the germline cells. As such, Drosophila species express piRNAs as well as piRNA pathway proteins in both the somatic and germline cells of the ovaries^7^.

piRNAs are abundantly produced from discrete genomic loci called piRNA clusters. In Drosophila ovaries, germline clusters are comprised of TE insertions in both orientations and produce piRNAs from both strands by virtue of non-canonical convergent transcription^8, 9^. In contrast, the somatic cluster *flamenco* in *D. melanogaster* consists of inverted repeats of *gypsy* retrotransposons and is transcribed from a canonical RNA Polymerase II promoter and predominantly produces *gypsy* antisense piRNAs^10, 11^.

There are two distinct modes of piRNA biogenesis, called “ping-pong” and “phasing”. Ping-pong biogenesis starts with the cleavage (termed slicing) of a precursor transcript by a pi-RISC which generates the 5’ end of a secondary piRNA. The secondary pi-RISC in turn cleaves a transcript from the opposite strand to generate another piRNA thereby amplifying both sense and antisense piRNA pools^10, 12^. Phasing biogenesis starts with 5’ end of a piRNA precursor transcript to be loaded onto a PIWI protein, which is followed by the head to tail fragmentation into mature piRNAs by the mitochondrial endonuclease Zucchini/MitoPLD^13, 14^. Both ping-pong and phasing mechanisms are evolutionarily highly conserved, present in sponge and hydra species all the way to humans^15, 16^.

Drosophila express three PIWI proteins in the ovaries, Piwi, Aubergine and Ago3. The somatic niche only expresses Piwi while the germline cells express all three^10^. Ping-pong biogenesis predominantly occurs between Aubergine and Ago3, while Piwi and Aubergine can receive phasing piRNAs. Ping-pong biogenesis is, therefore, specific to the germline, while phasing can happen in either cell niche.

For both, ping-pong and phasing, a mechanism to select and trigger piRNA biogenesis is necessary to specifically produce TE antisense piRNAs. The ping-pong mechanism itself provides the solution via the production and binding of sense piRNAs, which target and enrich for TE antisense piRNAs via sequence complementarity. Phasing, however, requires a ‘triggering’ event instead^17^. One way to trigger phasing in the Drosophila ovaries, is via slicing. The same pi-RISC that is produced in the ping-pong cycle initiates phasing by cleaving a transcript and releasing the 5’ end of a piRNA precursor. This slicer activity has been demonstrated to be the main source of triggering phasing in the Drosophila germline^18^. This mechanism of triggering appears to be evolutionarily conserved, as mouse pachytene piRNAs also require slicer activity to start phased biogenesis^19, 20^.

In the Drosophila soma, where ping-pong is absent, the recruitment of Zucchini by the TDRD12 homolog Yb mediates phasing instead. Yb forms an organelle called Yb body at the nuclear periphery to recruit piRNA biogenesis factors, such as the RNA helicase Armitage, to the *flamenco*-derived piRNA precursor transcripts^21, 22^. While Armitage is sufficient to induce piRNA biogenesis independently^23^, Yb is required for the preferential production of piRNAs from the *flamenco* locus. Drosophila mutants for *yb* increases promiscuous piRNA production, which upregulates transposons and leads to female infertility^22, 24, 25^.

Given that these mechanisms ensure the production of TE antisense piRNAs, the proteins associated with these processes are likely highly conserved^26^. Indeed, the three PIWI proteins, Yb and Armitage are also present in ancient Drosophila species such as *D. virilis* and *D. mojavensis*.

However, our inspection on FlyBase (https://flybase.org/) failed to identify Yb homologs in the *obscura* group within Drosophila, as well as *D. eugracilis*. More strikingly, we also could not find a homolog of Ago3 in *D. eugracilis*. We characterised ovarian piRNA populations in these species, finding that the *obscura* group as well as *D. eugracilis* predominantly produce TE antisense piRNAs in the soma despite the lack of Yb. We further find that ping-pong biogenesis is entirely absent in *D. eugracilis*, and that TE antisense piRNAs are produced by phasing without slicing in the germline. The study therefore reveals novel routes by which TE antisense transcripts are selected for piRNA production.

## Results

### *yb* gene is lost in species of the *D. obscura* group and *D. eugracilis*

In *D. melanogaster*, the depletion of Yb disrupts the formation of the processing body and disperses other biogenesis factors in the cytoplasm^27, 28^ (Figure 1A). This results in a marked decrease (> 10-fold) in the level of *flamenco*-derived piRNAs and the overexpression of *gypsy* transposons by more than a hundred-fold^22, 24^ (Figure 1B).

**Figure 1.**
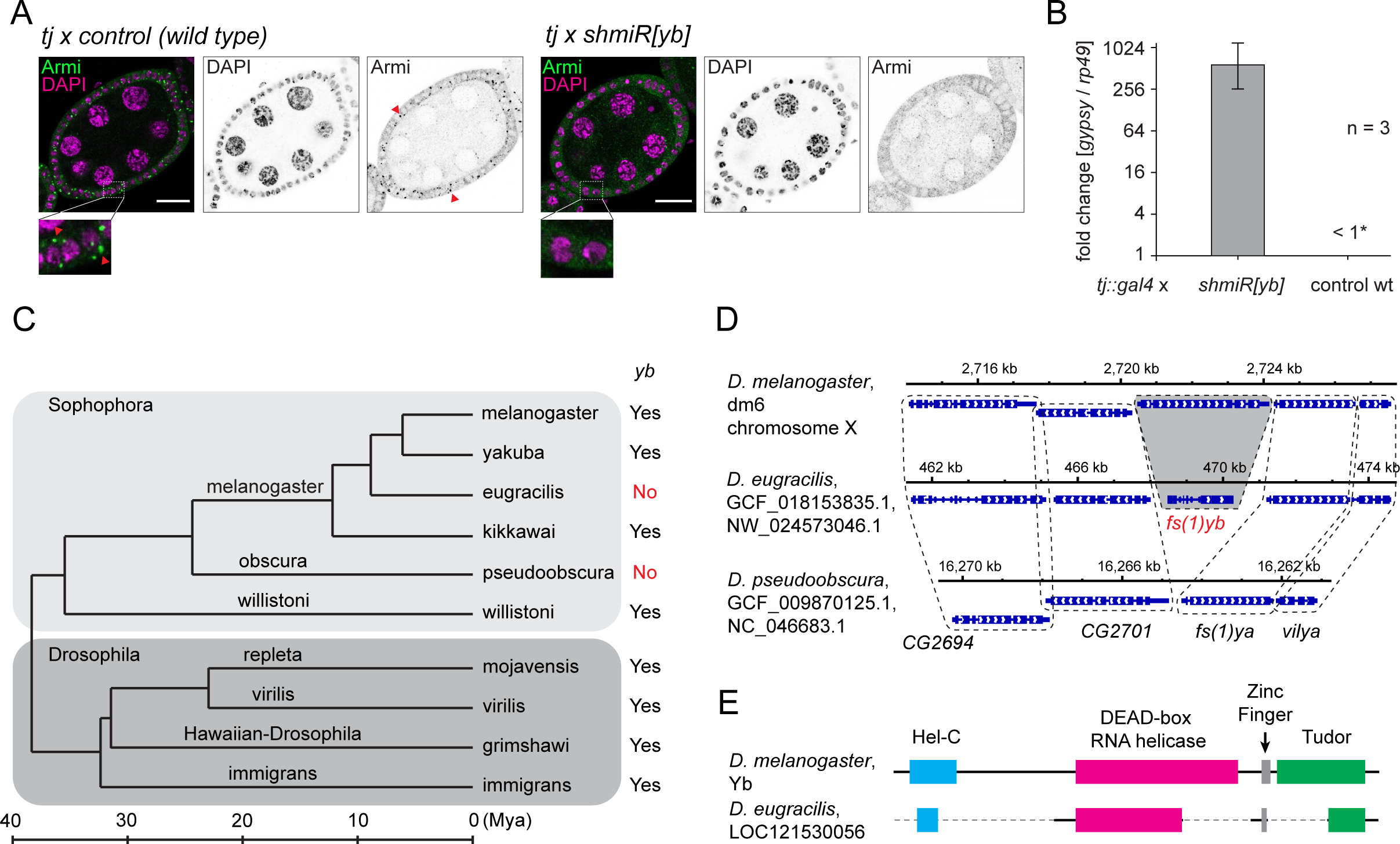
*fs(1)yb* (*yb*) is not conserved in the *obscura* group or in *D. eugracilis*. **(A)** Immuno-fluorescent staining of Armitage (Armi in green and DAPI in magenta) in the Yb-depleted (tj::gal4 x shmiR[yb]) and the wild type control (tj::gal4 x control) egg chambers shows that the focal localisation of Armi in the somatic cells (marked by read arrowheads) requires Yb. Enlarged images also highlight the Armi foci in the wild type somatic cells. Scale bars = 20 μm (**B)** An RT-qPCR shows that the depletion of Yb in the somatic cells results in an overexpression of *gypsy* (see methods). (**C)** The phylogenetic tree of selected Drosophila species showing the conservation of *yb* gene. The subgenera *Sophophora* and *Drosophila* are grouped in boxes and species groups are indicated on branches. Years of divergence and distances between species are based on previous studies ^58, 59^. (**D)** The conservation of the neighbouring genes in the syntenic locus of *yb* shows the degeneration and loss of *yb* gene in *D. eugracilis* and *D. pseudoobscura*, respectively. (**E)** The degeneration of *yb* gene in *D. eugracilis* is shown alongside with the functional domains.

Yb is encoded by the *female sterile 1 yb* (*fs(1)yb*) gene (named *yb* gene hereafter) in *D. melanogaster*. Unexpectedly, FlyBase did not annotate the *yb* gene in some of the most commonly studied Drosophila species, including *D. pseudoobscura* and *D. persimilis* from the *obscura* group. To extend this observation, we examined the conservation of *yb* in all Drosophila species whose genome sequences were available in high quality (see methods). This survey revealed that nearly all Drosophila species outside the *obscura* group species carry homologs of *yb* (Figure 1C, Supplementary Table 1). Importantly, the N-terminally-located Hel-C domain was found in all *yb* homologs, suggesting the functional conservation of *yb* gene across Drosophila species. Strikingly, none of the genomes from the twelve *obscura* group species analysed contained *yb* homologs whereas homologs of *boyb* and/or *soyb* were found, demonstrating the lack of *yb* gene in the *obscura* group species (Supplementary Table 1). Of Drosophila species outside the *obscura* group, we also failed to find the *yb* homologs in three independent genome assemblies of *D. eugracilis*. Crucially, flanking genes of *yb* are present at the syntenic locus of the *D. pseudoobscura* and *D. eugracilis* genomes (Figure 1D). These genes are also found next to the *yb* homologs of more ancient Drosophila species *D. willistoni* and *D. virilis* (data not shown), indicating that the *yb* gene has been specifically lost in these species. We also found that the fragments of domains in Yb are still present in the *D. eugracilis* genome (Figure 1E), but no mRNA expression for these fragments was detected by RNA sequencings in the ovaries (Figure S1), indicating it has recently become a pseudogene. The losses of *yb* in the *obscura* group and *D. eugracilis* likely occurred independently, as species which are evolutionarily closer to *D. eugracilis* than the *obscura* group species, have intact *yb* genes (Figure 1C).

### *gypsy-env* retrotransposons are present in species that have lost *yb*

The absence of *yb* in the *obscura* group and *D. eugracilis* prompted us to search for full-length copies of *gypsy* retrotransposons that contain *env* genes (*gypsy-*env) which are normally silenced by Yb-dependant piRNAs. Using existing annotations on RepBase, we identified several *gypsy* elements that carry sequences homologous to *env,* in *D. pseudoobscura*, *D. eugracilis*, as well as two other *obscura* group species; *D. bifasciata* and *D. azteca* (see methods). We also observed that each identified *gypsy-env* had multiple intact copies in their genomes (Supplementary Tables 2 and 3). These observations strongly suggest that active copies of the *gypsy-env* retrotransposons remain in the *obscura* group species and in *D. eugracilis,* despite the absence of *yb*.

### Both germline and somatic PIWI are present in the *obscura* group species and *D. eugracilis*

The lack of *yb* also prompted us to examine the expression of the PIWI proteins. A tBlastn search confirmed that all three PIWI genes, *piwi*, *aubergine*, and *Ago3* are present in the *obscura* group species, with *aubergine* being duplicated or triplicated in some of them (summarised in the Supplementary Table 4). However, a tBlastn search of Ago3 homologs identified the homolog of Aubergine as the top hit in the three independently assembled *D. eugracilis* genomes, suggesting the absence of the *Ago3* gene in this species. The consequences of the apparent loss of *Ago3* will be described in the later part of this work. We then analysed the localizations of Piwi, Aubergine, and Ago3 in *D. pseudoobscura* and *D. eugracilis* ovaries with antibodies raised against Piwi, Aubergine, and Ago3 (see methods). Piwi was nuclear both in the somatic and germline cells. Aubergine and Ago3 were only expressed in the germline and localised to the nuclear periphery. Ago3 also formed large granules near the nurse cell nuclei in *D. pseudoobscura* ovaries (Figure 2A) as well as in other *obscura* group species (Figure S2C). Additionally, Aubergine and Piwi localise to the pole cells and other somatic nuclei in the *D. eugracilis* blastoderm-stage embryos (Figure S2D). These observations suggest that both the germline and the somatic piRNA pathways are active in the *obscura* group species and *D. eugracilis,* and their PIWI proteins function similarly to those in *D. melanogaster*.

**Figure 2.**
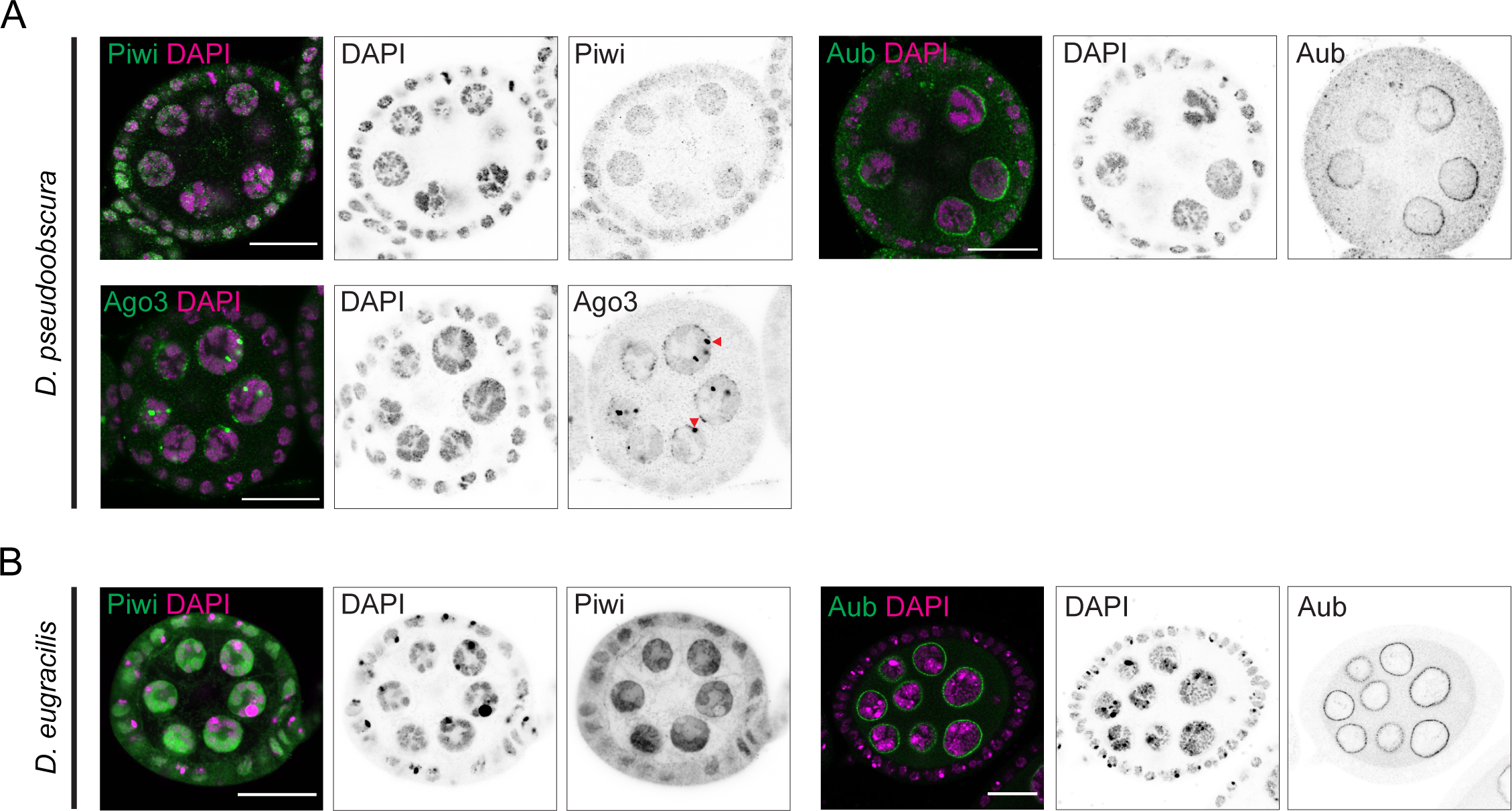
Conserved localisation of PIWI proteins in *D. pseudoobscura* and *D. eugracilis* ovaries. **(A)** Immuno-fluorescent staining of Piwi, Aubergine, and Ago3 in *D. pseudoobscura* egg chambers (antibodies in green and DAPI in magenta) showing the nuclear localisation of Piwi in both somatic and germline cells, and the peri-nuclear localisation of Aubergine and Ago3 only in the germline cells. Large peri-nuclear granules of Ago3 are indicated by arrowheads. **(B)** Immuno-fluorescent staining of Piwi and Aubergine in *D. eugracilis* egg chambers (antibodies in green and DAPI in magenta) showing the nuclear localisation of Piwi in both somatic and germline cells and the peri-nuclear localisation of Aubergine. Scale bars = 20 μm

### *flamenco*-like uni-strand clusters produce abundant piRNAs in the *obscura* group species

The *flamenco* locus in *D. melanogaster* predominantly consists of inverted repeats of *gypsy* retrotransposons. We investigated whether similar loci exist in species that do not have *yb*, and if so whether they make abundant piRNAs in the soma. We sequenced oxidised small RNA libraries from whole ovaries of the two *obscura* group species *D. pseudoobscura* and *D. bifasciata*. Inspections of small RNAs that uniquely mapped to the genome coupled with transposons predicted by RepeatMasker, identified *flamenco*-like piRNA clusters in these species (Figure 3A, Figures S3A and B). Importantly, the *gypsy-env* retrotransposons that we identified in these species have insertions in the clusters (indicated by colours in the figures), suggesting their role in suppressing them. We also found clusters that resemble germline clusters in *D. melanogaster,* in that they produce abundant piRNAs from both strands and have little bias in the orientation of *gypsy* insertions (Figure 3B and Figure S3C). We performed fluorescent in-situ hybridisation (FISH) using short-oligo DNA probes to examine the expression of the precursor transcripts from the clusters in the ovaries. FISH signals of the uni-stranded clusters were detected in the somatic cells, while the dual-stranded clusters were expressed in the germline cells in *D. pseudoobscura* and *D. bifasciata* (Figures 3C – D and Figures S3D - F). We further observed that these uni-stranded clusters have putative promoter peaks of RNA polymerase II and spliced precursor transcripts, resembling protein coding genes (Figures S4A-C). These are features common to the *flamenco* cluster in *D. melanogaster*^8^. We conclude that *D. pseudoobscura* and *D. bifasciata* have both somatic and germline piRNA clusters that resemble those of *D. melanogaster*.

**Figure 3.**
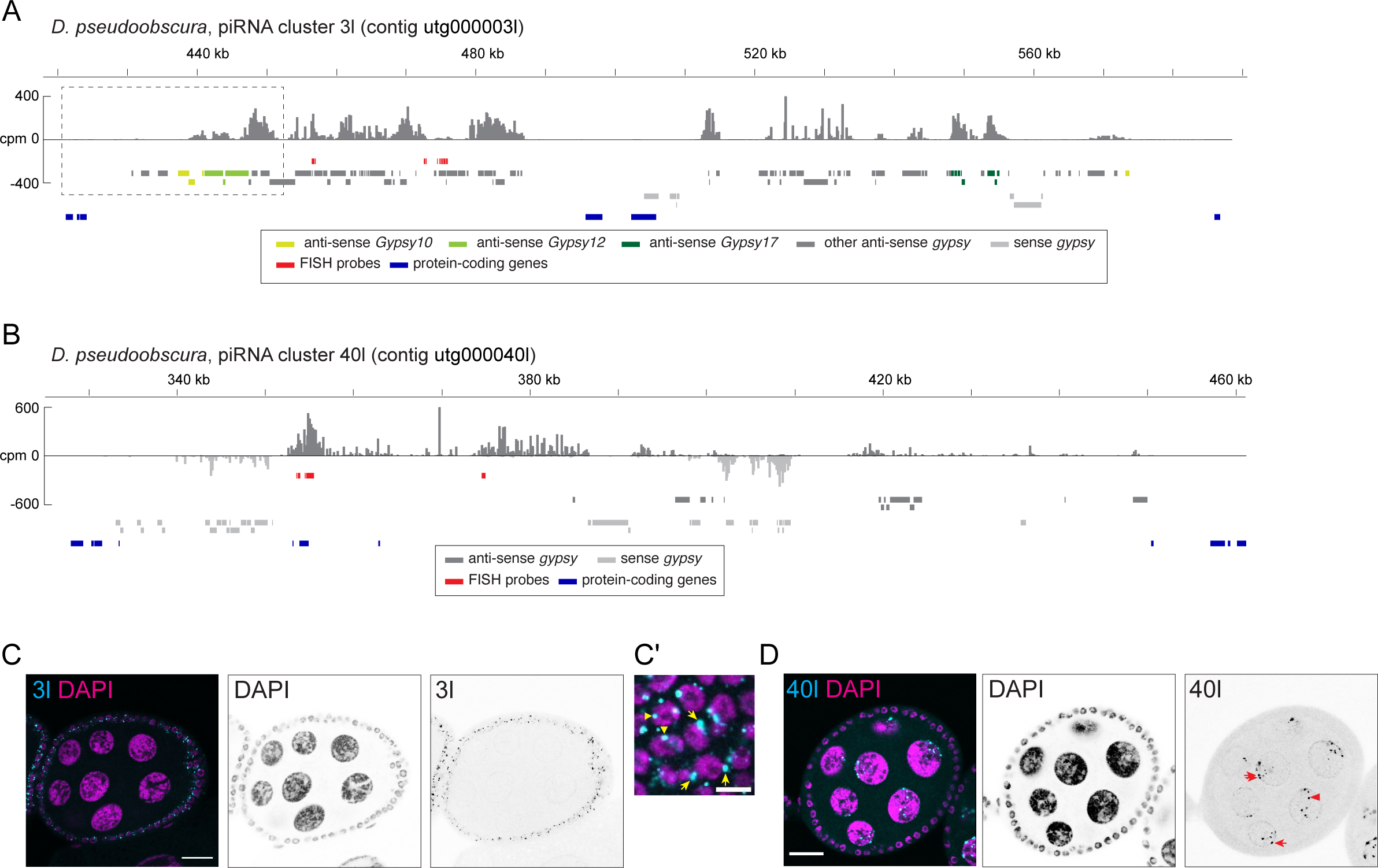
Somatic and germline piRNA clusters in *D. pseudoobscura* ovaries resemble those of *D. melanogaster*. **(A) and (B)** Shown is the coverage of piRNA reads (>22nt) in counts per million genome mappers (CPM) from the oxidised whole ovary small RNA library of *D. pseudoobscura* that uniquely mapped to the cluster regions. Sense and antisense reads are coloured in dark and light gray, respectively. Coloured bars indicate *gypsy* insertions predicted by RepeatMasker, annotated protein-coding mRNA exons and the FISH probes. A dotted box in (A) indicates the putative transcription start site of the cluster, for which a magnified view is shown in Figure S4A. **(C) and (D)** RNA FISH against transcripts from the piRNA clusters, showing the expression of the cluster 3l (C and C’) and 40l (D) in the somatic and germline cells, respectively. A birds’ eye-view of the somatic epithelium of a FISH-staining of 3l is shown in (C’). Putative sites of transcription (in the nuclei) and processing (at the nuclear periphery) of the cluster transcripts are indicated by arrowheads, and arrows, respectively (C’ and D). Scale bars = 20 μm in (C) and (D) and 5 μm in (C’).

We also found uni-stranded and dual-stranded piRNA clusters in *D. eugracilis* with similar arrangements of *gypsy* insertions (Figure S5). We could not determine the cell-type specific expression of *D. eugracilis* clusters by FISH as they expressed much fewer piRNAs per kilo bases than clusters from the other species (data not shown). The uni-stranded cluster that we named 3031 produces piRNAs antisense to copies of *env*-carrying *Gypsy-6_Deu*, suggesting its role in silencing the element in the soma, and resides within an intron of a protein-coding gene (Figure S5C).

### Somatic biogenesis bodies are present in the *obscura* group species and *D. eugracilis*

piRNA biogenesis factors concentrate on cytoplasmic *flamenco* RNA to form Yb-bodies in *D. melanogaster*. In the absence of Yb, other biogenesis factors, such as Armitage, disperse in the cytoplasm and fail to produce abundant piRNAs from *flamenco*^24^ (Figure 1A). The immuno-fluorescence staining in *D. bifasciata*, *D. subobscura*, *D. azteca*, and *D. eugracilis* ovaries showed focused localisation of Armitage in the somatic cells, resembling the *D. melanogaster* Yb body (Figures 4A-D). Furthermore, a co-staining of Armitage and the transcripts from the uni-stranded cluster CM1_137 identified earlier in *D. bifasciata* ovaries revealed their co-localisation in the 3D-reconstructed images (Figure 4E), indicating the presence of the piRNA biogenesis bodies despite the absence of Yb.

**Figure 4.**
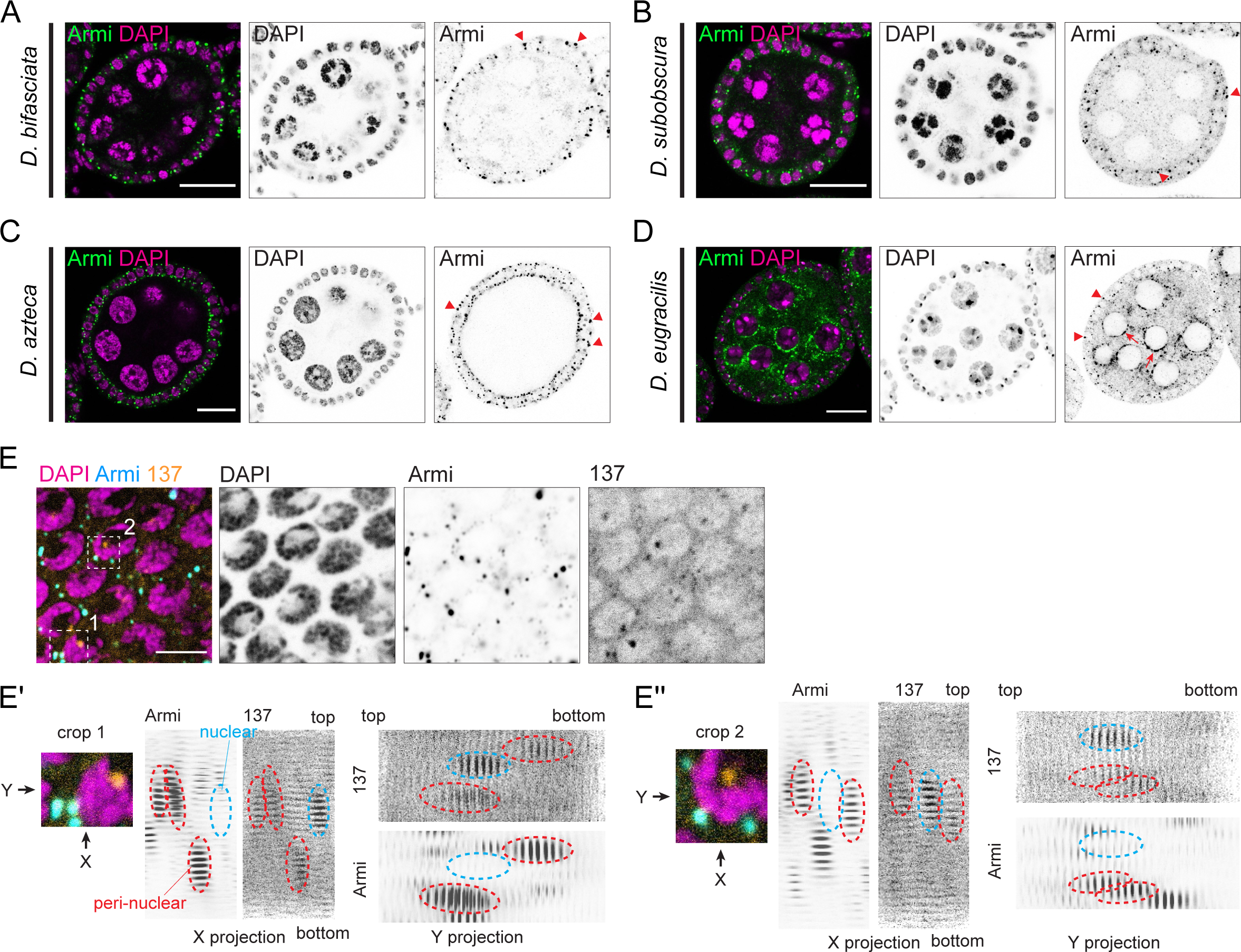
Somatic piRNA biogenesis bodies in the absence of Yb. **(A) - (D)** Immuno-fluorescent staining of Armitage in *D. bifasciata* (A)*, D. subobscura* (B)*, D. azteca* (C)*, and D. eugracilis* (D) egg chambers (Armitage in green and DAPI in magenta) showing the focused localisation in the somatic epithelium (arrowheads) and the peri-nuclear localisation in the germline (arrows). **(E)** Co-staining of Armitage (blue) and the transcript RNA from the cluster CM1_137 (orange) in a somatic epithelium of a *D. bifasciata* egg chamber. **(E’) and (E**’’) Shown are 3D-reconstructions of Z-stack images taken at regions indicated in (E). About 30 slices of confocal planes from the Armitage and the cluster CM1_137 stainings are aligned and projected from the X and Y axes to observe the co-localisation. Nuclear and peri-nuclear signals of CM1_137 are indicated by cyan and red dotted circles, respectively. Scale bars = 20 μm in (A) - (D) and 5 μm in (E).

### Specialised somatic piRNA biogenesis for *gypsy* is evolutionarily conserved in *Drosophila*

The focal localisation of Armitage in the *obscura* group species and *D. eugracilis* indicates that they have still have mechanisms to efficiently process the cluster-derived transcripts into piRNAs. To further characterise somatic piRNAs, we sequenced the embryonic (0 – 2 hr post fertilisation) small RNA pool, which is devoid of somatic material, and compared it to whole ovarian small RNAs.

We first confirmed that the method faithfully captures piRNAs from somatic and germline compartments. Firstly, piRNAs targeting known somatic retrotransposons were scarce in the embryonic pool in *D. melanogaster*^7, 29^ (Figure S6A). Furthermore, the piRNAs mapping to the somatic *flamenco* locus were enriched by more than 10-fold in the ovarian pool, and were amongst the most abundant piRNAs in the soma. In contrast, the germline cluster 80F was equally represented in both libraries (Figure 5B). We further observed that about three quarters of piRNAs enriched in the ovaries come from discrete clusters like *flamenco*, or from other *gypsy* antisense insertions (Figure 5E). We finally estimated the abundance and composition of ovary-enriched somatic piRNAs in Yb-depleted *D. melanogaster* flies. We observed a reduction in *flamenco*-derived piRNAs by around 15-fold in the ovaries, while genic piRNAs were reduced by only 18% (Figure 5E and Supplementary Table 5). In contrast, the control RNAi ovaries expressed comparable levels of *flamenco*-derived piRNAs alongside other *gypsy* antisense piRNAs which corroborates with previous findings^24^.

**Figure 5.**
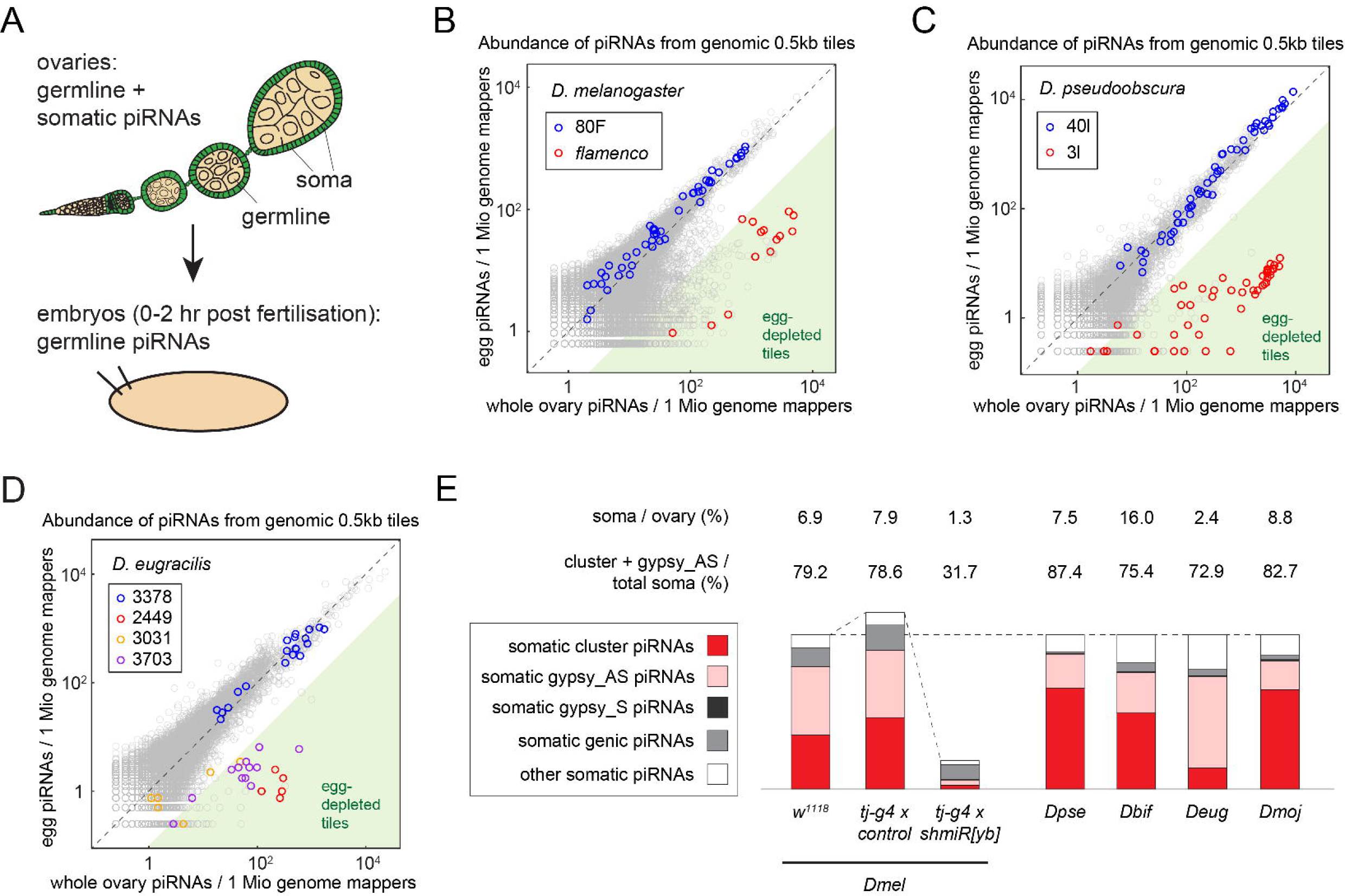
Somatic piRNA biogenesis in *Drosophila* species is highly specialised for targeting *gypsy*. **(A)** A schematic illustration contrasting the whole ovaries and the embryos that consist of both somatic (green) and germline (beige) cells, and germline cells alone, respectively. Somatic piRNAs are only represented in the whole ovary small RNA pool while germline piRNAs are represented in both libraries. **(B) - (D)** Scatter plots showing the abundance of piRNAs from the whole ovaries (X axes) and the eggs (Y axes) that uniquely mapped to the individual 0.5 kb tiles of *D. melanogaster* (B), *D. pseudoobscura* (C) and *D. eugracilis* (D) genomes. Dual-stranded germline clusters are coloured in blue while uni-stranded somatic clusters are coloured in orange, red and purple. Tiles that expressed piRNAs more than ten times in the whole ovaries than in the eggs are shaded in green. **(E)** Bar charts showing the abundance of different classes of somatic piRNAs from indicated genotypes and species. The estimated proportions of somatic piRNAs per ovarian piRNAs, the sum of cluster-derived piRNAs and *gypsy* antisense piRNAs per total somatic piRNAs are shown. The bars are adjusted to the same height for the wild type strains and are relative to the abundance of somatic piRNAs for the three strains of *D. melanogaster*. Species names are abbreviated as follows: *D. melanogaster*; Dmel, *D. pseudoobscura*; Dpse, *D. bifasciata*; Dbif, *D. eugracilis*; Deug, and *D. mojavensis*; Dmoj.

The comparison of the embryonic and ovarian piRNA pools in the *obscura* group species and *D. eugracilis* mirrored the observations in *D. melanogaster*. Firstly, the transposons that abundantly produce somatic piRNAs are classified in the same subgroup as *gypsy*, indicating that somatic piRNAs in these species defend against similar types of transposons as *D. melanogaster*^29^ (Figures S6B and C). Secondly, piRNAs mapping to the somatic clusters identified earlier were enriched in the ovaries, while the germline clusters were equally represented in both libraires (Figures 5C and Figure S6D). Additionally, we found that other uni-stranded clusters in *D. bifasciata* and *D. eugracilis* produce piRNAs in the soma. We further found that piRNAs from discrete clusters and from *gypsy* antisense insertions constitute a majority of somatic piRNAs (Figure 5E, 87.4%, 75.4%, and 72.5% for *D. pseudoobscura*, *D. bifasciata*, and *D. eugracilis*, respectively). The numbers for *D. bifasciata* and *D. eugracilis* were likely underestimated due to relatively poor annotations of transposons in these genomes. These observations indicate that the somatic piRNA pathway in these species remains highly selective, despite the lack of Yb.

Similar observations were also made in *D. mojavensis*, an evolutionarily distant species from the *Drosophila/Drosophila* subfamily (Figure 1). We found a *flamenco*-like uni-strand piRNA cluster and confirmed that the cluster produces abundant piRNAs in the soma (Figure S6). We also found that a majority of somatic piRNAs (82.7%) are either from discrete clusters or *gypsy* antisense insertions (Figure 5E). These observations indicate that specialised somatic piRNA biogenesis for *gypsy* appears to be a universal feature in the *Drosophila* taxa, regardless of the conservation of *yb*.

### Absence of ping-pong piRNAs in *D. eugracilis*

Ago3 is primarily required to bind transposon sense piRNAs to maintain the ping-pong amplification loop in *D. melanogaster*. Despite the absence of *Ago3* in addition to *Yb*, *D. eugracilis* ovaries still produce piRNAs as abundantly as other species (Figure 6C). Strikingly, when we further examined these piRNAs, we found that most transposon piRNAs originate from the antisense strand. Of the 84 most piRNA-producing transposons in the *D. eugraclis* germline, 67 of them showed more than 95% bias towards antisense (Figure 6B). For example, the piRNAs mapping to *BEL-2_Deu* are almost exclusively antisense (Figure 6A). In contrast, similar transposons in *D. melanogaster* and *D. pseudoobscura* produce 15-40% of sense piRNAs (Figure S7). The observed absence of sense piRNAs prompted an examination of ping-pong in *D. eugracilis*. We measured the distance between the 5’ ends of overlapping transposon sense and antisense piRNAs, where ping-pong pairs show a characteristic 10nt overlap. Unexpectedly, of the ten germline *D. eugraclis* transposons that produce at least 10% of sense and antisense piRNAs, none demonstrated any enrichment at the 10nt overlap, highly suggestive of a lack of ping-pong in *D. eugracilis* (Figure 6D and Figure S7C). In contrast, most germline transposons in *D. melanogaster* and *D. pseudoobscura* show strong tendencies of having the 10nt overlap to the antisense piRNAs (Figure 6D, examples shown in Figure S7).

**Figure 6.**
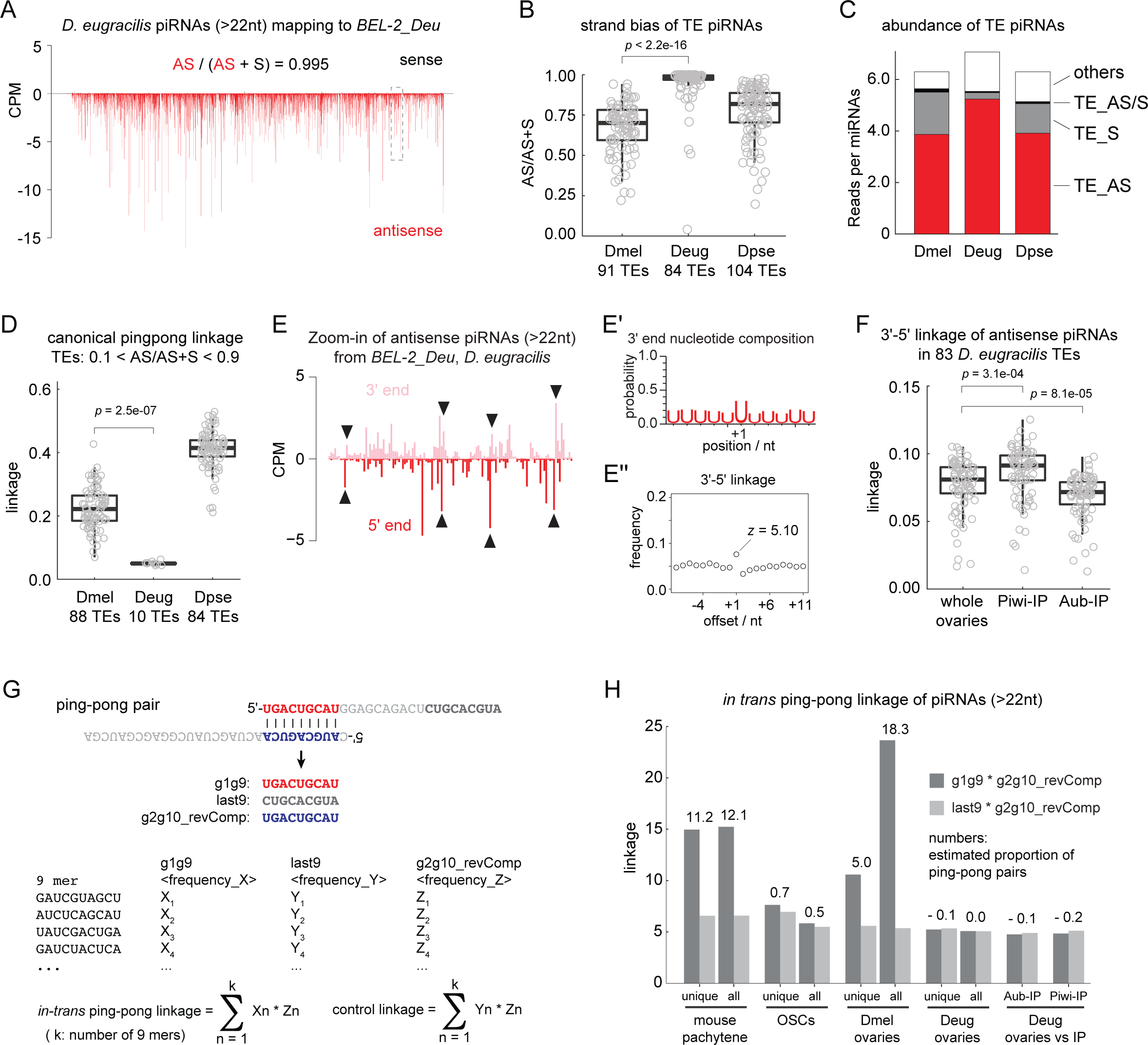
Slicer-independent phasing predominantly produces transposon antisense piRNAs in *D. eugracilis* germline. **(A)** Shown are the 5’ end coverage of piRNA reads (>22nt) mapping to *BEL-2_Deu* from the oxidised whole ovary small RNA library of *D. eugracilis* in counts per million genome mappers (CPM). Sense and antisense reads are coloured in black and red, respectively. Antisense (AS) piRNAs make up more than 99% of all piRNAs mapping to *BEL-2_Deu*. **(B)** A box plot showing the strand bias of transposon-mapping piRNAs measured by the fraction of antisense reads out of the total reads. Circles represent individual transposons. Top ∼100 transposons that produce most piRNAs in respective species are shown. *p-value* is calculated by Mann-Whitney U test. **(C)** A bar chart showing the abundance of piRNAs of different classes relative to the abundance of microRNAs. Transposon piRNAs are grouped into sense (TE_S), antisense (TE_AS) and both (TE_AS/S). **(D)** A box plot showing the canonical ping-pong linkage of transposon-mapping piRNAs. Only the transposons that express at least 10% of both sense and antisense piRNAs are included. *p-value* is calculated by Mann-Whitney U test. **(E)** Shown are the 5’ and 3’ ends of antisense piRNAs mapping to *BEL-2_Deu* from the dashed box in (A). piRNA 5’ ends are frequently found one nucleotide downstream of piRNA 3’ ends as indicated by arrowheads. **(E’)** Shown are the frequencies of Uridines found at positions relative to the 3’ ends of piRNAs mapping to *BEL-2_Deu*. +1 corresponds to the immediate downstream nucleotide position. **(E’’)** Shown is the frequency plot of the 3’-5’ linkage of antisense *BEL-2_Deu* piRNAs. The z score of the linkage position +1 is shown (see method). **(F)** A bar chart showing the 3’-5’ linkage values of transposon antisense piRNAs comparing the total ovarian piRNAs to Piwi- and Aubergine-bound piRNAs. *p-values* are calculated by Mann-Whitney U test. **(G)** A summary of *in-trans* ping-pong analysis. 9mers from positions g1 to g9 (g1g9), last nine nucleotides (last9) and the reverse complemented g2 to g10 (g2g10_revComp) are extracted from all piRNA reads and frequencies of individual 9mer sequences are measured for each group. *in-trans* ping-pong linkage is calculated as the sum-product of frequencies between g1g9 and g2g10_revComp found in the same sequences while the sum-product between last9 and g2g10_recComp serves as a negative control. **(H)** Shown are *in-trans* ping-pong linkage values of genome unique piRNAs and all piRNA mappers per samples. Estimated proportions of *in-trans* ping-pong pairs out of all piRNAs are shown in percentage. “g2g10_revComp” of Aubergine-(Aub-) and Piwi-bound piRNAs were compared against “g1g9” and “last9” of the total ovarian piRNAs where indicated.

### Piwi and Aubergine receive phasing piRNAs in *D. eugracilis*

Although *D. eugracilis* piRNAs lack ping-pong signatures, they retain other characteristics of piRNAs seen in *D. melanogaster,* such as their size and preference to start with a Uridine at the 5’ end (Figure S8B). During phased piRNA biogenesis, the endonuclease Zucchini/MitoPLD preferably cleaves in front of a Uridine simultaneously produce the 3’ end of the preceding and the 5’ end of the next piRNAs. Transposon piRNAs in *D. eugracilis*, such as those mapping to *BEL-2_Deu*, demonstrate characteristics associated with phasing including the characteristic head-to-tail arrangement of flanking piRNAs, as well as a Uridine bias immediately downstream of the 3’ end of the piRNAs (Figure 6E).

We further explored this preference to phasing biogenesis by performing an immuno-precipitation using specific antibodies for *D. eugracilis* Piwi and Aubergine, sequencing the small RNA pools and validating them against the embryonic small RNA sequencing (Figures S8A and G). Piwi, Aubergine, and Ago3 in *D. melanogaster* participate in ping-pong and phasing to varying extents and receive different pools of piRNAs. In contrast, we found that Aubergine and Piwi receive nearly identical populations of piRNAs in *D. eugracilis*, as seen in the size distribution, sense-antisense bias of transposon piRNAs and at the level of individual piRNAs (Figures S8C to F). In addition, both proteins receive phasing piRNAs as indicated by the 3’-5’ linkage of the transposon antisense piRNAs (Figures S8E and F, Figure 6F). These observations strongly indicate that *D. eugracilis* predominantly rely on phasing for producing piRNAs in the germline.

### Phasing occurs without slicing by PIWI proteins in the *D. eugracilis* germline

Abundant phasing in the absence of ping-pong piRNAs in the *D. eugracilis* germline was not expected, as the slicing event by a PIWI RISC, as part of the ping-pong cycle, is required to trigger phasing from the RNA substrate in the *D. melanogaster* germline. On the other hand, slicing can occur without perfect sequence complementarity of the entire length of piRNA. Therefore, a piRNA produced from one genomic locus can initiate phasing from different loci *in-trans*, such as in mouse pachytene spermatocytes^20^ (Figure S9A). To test whether germline piRNAs in *D. eugracilis* are produced in a similar manner, we devised a computational tool, which we named “*in-trans* ping-pong analysis” to measure the frequency of genome-wide slicing-triggered phasing events. Although the precise rule of piRNA target recognition is incompletely understood, the region between g2 to g10 positions is minimally expected to match the target sequence^30, 31^. We counted the frequencies of every 9mer found at the g1-g9 part of piRNAs and measured how often they find identical sequences at the reverse-complemented g2-g10 of other piRNAs (see methods and Figure 6G). Consistent with the previous findings^19, 20^, mouse pachytene-stage piRNAs showed a strong signature of *in-trans* ping-pong where more than 10% of piRNAs were estimated to have pairs (Figure 6H). As a negative control, fewer than 1% of piRNAs from the ovarian somatic cell line (OSCs, *D*. *melanogaster*), which is devoid of slicing-competent PIWI proteins, could find the ping-pong pairs. The *D. melanogaster* whole ovarian piRNAs showed strong *in-trans* ping-pong linkage, which also measures the abundance of canonical ping-pong pairs. The analysis estimated that there were fewer ping-pong pairs between uniquely mapped piRNAs (5.0%) than all piRNAs mappers (18.3%) in *D. melanogaster*. This is also seen in *D. pseudoobscura* piRNAs and consistent with the abundance of ping-pong piRNAs from transposons (Figure 6H and Figure S9C). Strikingly, *in-trans* ping-pong pairs were not detected in *D. eugracilis* either in the pool of all genome mapping piRNAs or in the pool of genome-unique mappers (Figure 6H and Figure S9C). An underrepresentation of Piwi-or Aubergine-bound piRNAs, which may take part in slicing-triggered phasing but make up a small fraction of the total piRNA pool, can be ruled out. This is because neither of them alone showed any linkage of *in-trans* ping-pong to the total pool of piRNAs (Figure 6H). We conclude that phasing piRNA biogenesis occurs independently of slicing in the germline of *D. eugracilis*.

## Discussion

We analysed Drosophila species that lack key known piRNA biogenesis factors in *D. melanogaster*. We demonstrated that the *obscura* group species and *D. eugracilis* are able to abundantly produce somatic *gypsy* antisense piRNAs in the absence of *yb*. We further demonstrated that *D. eugracilis* exclusively produces phasing piRNAs in the germline, independently of slicer activity.

### Somatic cluster Architecture is conserved across Drosophila

piRNA clusters are highly diverged in their genomic origins but retain general structural features over evolution^32–34^. A recent study showed that individual germline clusters are dispensable for silencing transposons in *D. melanogaster*, suggesting a rapid change in transposon contents and a slower turn-over of cluster identities over evolution^35^. In contrast, the *flamenco* cluster in *D. melanogaster* is required for suppressing *gypsy* and maintaining female fertility. Here, we demonstrated that the *flamenco*-like cluster appears to be deeply conserved in Drosophila, which we characterised by its strand bias, *gypsy* insertions, and somatic cell specificity. These clusters also carry copies of active *gypsy*, suggesting their functional importance. It is important to note that *env*-carrying *gypsy* elements are not only evolving within species, but also frequently transmitting between species likely due to their capacity to form virus-like particles^29^. It is therefore likely that the somatic piRNA clusters are under greater pressure than the germline clusters to adapt to rapidly changing *gypsy* elements.

### Gypsy antisense piRNAs can be produced independently of Yb

Despite the loss of *yb*, the *obscura* group and *D. eugracilis* are able to produce *gypsy*-antisense piRNAs to a high degree of specificity. The biogenesis factor Armi appears to form Yb-body-like structures that co-localise with cluster-derived RNA in these species, which indicates that the downstream processing of piRNAs occurs in a similar manner as *D. melanogaster*. One possibility is that another protein has taken over the role of Yb by recruiting Armitage to the cluster transcripts. Since neo-functionalisation of duplicated gene paralogs is frequently found in the piRNA pathway^9, 36, 37^, other Tudor proteins or DEAD-box RNA helicases may have once again duplicated in these species to replace Yb. Alternatively, the clusters in Yb-less species might be transcriptionally regulated in a way that transcripts are efficiently processed by Armitage in the absence of Yb. This idea seems consistent with the high ratio of cluster/TE piRNAs to genic piRNAs observed in this study (Figure 5H).

### Only ping-pong/ No slicer activity can produce Phased piRNAs

A lack of Ago3 in *D. melanogaster* leads to the sense and antisense piRNAs being loaded onto Aubergine for a homotypic form of ping-pong^38, 39^. Though sterile, the absence of Ago3 does not erase ping-pong signature in the ovaries. In contrast, Aubergine in *D. eugracilis* exclusively receives phased piRNAs, and no ping-pong signature was observed.

Other species, such as *C. elegans*, have also diverged away from the standard piRNA pathway, and rely on other biogenesis pathways to specify TE piRNAs^40^. Curiously, *D. eugracilis* still appears to have much of the central mechanisms behind standard piRNA production, such as subcellular localisation and the 5’ Uridine preference of PIWI proteins. Furthermore, piRNA biogenesis factors that are known to be involved in ping-pong, such as Spindle-E^7^, Vasa^41^ and Qin^17^, are all conserved in *D. eugracilis*. Although the precise mechanism is not known, these factors bind and stabilise PIWI proteins with precursor RNA at different steps in the ping-pong cycle^41, 42^. Whether any of these factors participates in piRNA biogenesis in *D. eugracilis* and whether there is common logic in their molecular actions, are both intriguing open questions.

The lack of ping-pong in *D. eugracilis* may have broader implications on the piRNA pathway as a whole, such as how germline piRNA clusters are transcribed and processed, let alone the driving force to lose such a robust and evolutionarily deeply conserved mechanism^16^. The Rhino-Cutoff-Deadlock (RDC) complex in *D. melanogaster* binds germline piRNA clusters, and couples transcription to nuclear export, then to piRNA biogenesis^9, 43–45^. Most of the players in this circuit are again conserved at the gene level in *D. eugracilis*. However, this alone does not explain the extreme strand bias of transposon piRNAs in *D. eugracilis,* as Rhino-licensed transcription is bi-directional by nature utilising dispersed promoters within the clusters^36^. On the other hand, outside promoters also allow transcription through the cluster, potentially by suppressing splicing via the RDC complex^9, 36, 46^. Although the available genome assembly did not allow us to examine the possibility of outside promoters, *flamenco*-like clusters may exist in the *D. eugracilis* germline where transposons are aligned to the same direction and the strand bias of piRNAs is dictated by external promoters.

The lack of ping-pong, and the lack of piRNA-guided slicing as a whole, introduces another interesting conundrum. Phasing biogenesis requires a ‘triggering’ event to initiate the endonucleolytic cleavage of piRNA precursors (Figure S10). In the *D. melanogaster* germline, the slicing activity of Ago3-bound and Aubergine-bound piRNAs is required for the production of nearly all germline phased piRNAs bound by Piwi^17, 18^. It is possible that the *D. eugracilis* germline has become like the soma and has become capable of bringing precursor transcripts to Zucchini on the mitochondrial surface, without the need of endonucleolytic cleavage. However, this mode of piRNA biogenesis would be more promiscuous by nature in the absence of a “specificity” factor, such as Yb^24^. Therefore, *D. eugracilis* germline may have two distinct mechanisms to achieve specificity: the cluster definition and the selection of cluster-derived transcripts.

The logic of sequence complementarity of a piRNA and its target may go beyond ping-pong biogenesis. Maternally deposited Piwi- and Aubergine-bound piRNAs are responsible for silencing transposons in the next generation^47^. In transposon-induced hybrid dysgenesis, only the cross of a naïve father and a mother that carries loci that produce abundant transposon piRNAs can produce fertile progeny. It is thought that maternally inherited transposon piRNAs find the target RNA to produce more piRNAs from the same transposons. Therefore, the logic of sequence complementary is akin to the piRNA-dependent transgenerational silencing of transposons. We found that Aubergine and Piwi localise to the pole cells and other somatic nuclei in the *D. eugracilis* embryos before the zygotic transcription starts. Maternally inherited Aubergine and Piwi may serve the same purpose by other unknown mechanisms in *D. eugracilis*. Alternatively, they may have other functions, such as selectively degrading mRNAs in the germ plasm^48^ or suppressing transposons in the somatic cells during embryonic development^49^.

In summary, we found that the *obscura* group species and *D. eugracilis* have likely acquired novel mechanisms of phasing piRNA biogenesis distinct from those of *D. melanogaster* and other Drosophila species (summarised in Figure S10). This raises mechanistic questions of what is possible in the piRNA pathway, but also raises many biological conundrums to self and non-self RNA distinction.

## Materials and Methods

### Fly husbandry

We obtained wild type strains of *Drosophila pseudoobscura* (k-s12), *Drosophila eugracilis* (E-18102) and *Drosophila mojavensis* (k-s13) from the Drosophila species stock centre at the Kyorin University, *Drosophila bifasciata* (14012-0181-02), *Drosophila subobscura* (14011-0131-04), and *Drosophila azteca* (14012-0171-03) from The National Drosophila Species Stock Centre at the Cornell University. The *Drosophila melanogaster* strains, *w^1118^*, *armitage^1^* and *armitage^72.1^* were obtained from the Bloomington Drosophila Stock Centre. *traffic-jam gal4* and shmiR lines for *yb*^28^ and *armitage*^50^ were obtained from Dr Dorothea Godt, and Dr Julius Brennecke, respectively. Flies were raised at room temperature (∼22°C) in the standard food based on molasses, semolina, sugar, and fresh yeast.

### Identification of the *yb* homologues in *Drosophila* species

We examined the conservation of the *yb*, *boyb* and *soyb* genes in a total of 356 genome assemblies of Drosophila species that were available in NCBI in February 2022 as well as 15 Nanopore genome assemblies published in Miller et al^51^. We performed a tBlastn search using the default option of the stand-alone NCBI Blast package 2.9.0. The FlyBase entries FBpp0070462, FBpp0078210, and FBpp0292885 were used as the bait sequences for Yb, BoYb, and SoYb, respectively. The conservation of *yb* was called when the whole Hel-C domain (33-133aa) and the DEAD-box RNA helicase domain (391-740aa) found the homologous parts in a single locus. Additionally, the Hel-C domains of the Yb homologues from species that are distantly related to *D. melanogaster* were predicted by the HHpred (https://toolkit.tuebingen.mpg.de/tools/hhpred) and the whole protein sequences were used as baits for the tBlastn search. They include ALC48774.1 (*D. busckii*), XP_043072139.1 (*D. grimshawi*), XP_023179535 (*D. hydei*), XP_002055589.2 (*D. virilis*), XP_001965435 (*D. ananassae*), XP_017029052.1 (*D. kikkawai*), and XP_023034986.1 (*D. willistoni*). The results are summarised in Supplementary Table 1. We did not examine more than two genome assemblies from the same species when the *yb* homologues were identified in two independent assemblies.

### Identification of intact *env*-carrying *gypsy* insertions in *Drosophila* genomes

We ran RepeatMasker 4.1.0 to predict the *gypsy* insertions in the following genome assemblies: the UCI_Dpse_MV25 assembly (RefSeq accession: GCF_009870125.1) and the Nanopore assembly of the *D. pseudoobscura* genome (PMID: 30087105), the UCBerk_Dbif_1.0 assembly of the *D. bifasciata* genome, the DaztRS1 assembly (GenBank accession: GCA_005876895.1) of the *D. azteca* genome, and the ASM1815383v1assembly of the *D. eugracilis* genome. We used the RepBaseRepeatMaskerEdition-20181026 for finding transposable elements. Separately, we searched for the genomic loci that contain all Open Reading Frames of the *env*-carrying *gypsy* retrotransposons using the sequences obtained from the RepBase. We searched copies of *Gypsy10_DPse*, *Gypsy17_Dpse*, and *Gypsy12_Dpse* in the *D. pseudoobscura* genome, *Gypsy17_Dpse* and *Gypsy_DS* in the *D. bifasciata* genome, *Gypsy-3_DAzt*, *Gypsy-8_DAzt*, *Gypsy-19_DAzt*, and *Gypsy-101_DAzt* in the *D. azteca* genome, and *Gypsy1_DM*, *Gypsy-6_DEu* and *Gypsy-37_DEl* in the *D. eugracilis* genome. We called RepeatMasker-predicted *env*-carrying *gypsy* insertions as “intact” when all ORFs were present. The genomic loci of all identified copies and the detailed criteria of calling the intact insertions can be found in Supplementary Table 3.

### Immuno-fluorescence staining of ovaries

Rabbit polyclonal antibodies were generated by Genscript using the following peptides where additional single Cysteine residues for the cross-linking purpose are indicated in lowercase: *D. pseudoobscura* Piwi (MSENQGRGHRRPHGc), Aubergine (MNDLPTNSGHSRGRc), and Ago3 (MSGRGNLLKLFNKKc), *D. eugracilis* Piwi (GRRRPLYDEEPSTSc) and Aubergine (cANKQGDPRGPVSGR). Purified IgG were reconstituted in PBS (0.5 μg/ml) before use. All five antibodies predominantly detected single bands at expected sizes (100 ∼ 110 kDa) by western blotting (Figures S2A and B). Mouse monoclonal anti *D. melanogaster* Armitage (1D1-3H10), which was raised against the peptide from 36 - 230aa of FBpp0100102, is a gift from Julius Brennecke. The ovaries were freshly dissected from 2 – 7d old females in PBS and fixed in PBS containing 4% formaldehyde for 10 min at room temperature. Fixed ovaries were permeabilised in PBS containing 0.5% v/v Triton-X for 30 min, washed in PBS containing 0.1% Triton-X (PBS-Tx) several times before blocking in PBS-Tx containing 0.05% w/v BSA for 30 min. The primary antibody incubation was conducted in the blocking solution at 4°C overnight with antibodies in the following dilutions: 1:500 for anti *D. pseudoobscura* Piwi, Aubergine, and Ago3, anti *D-eugracilis* Piwi and Aubergine, and 1:200 for anti *D. melanogaster* Armitage. Goat anti rabbit or mouse IgG conjugated to Alexa Fluoro 488 or 568 (Abcam) were used as secondary antibodies and the confocal images were taken on a Zeiss LSM-800. DAPI was used to visualise DNA. Images were processed by Fiji.

### Immuno-fluorescence staining of embryos

*D. eugracilis* embryos of 1-1.5hr post fertilisation were collected and the chorions were removed by bleaching. Embryos were then fixed in heptane saturated by formalin for 30 min at room temperature, and washed by methanol to remove the vitelline membrane and the residual heptane. Subsequently, embryos were fixed again by PBS containing 4% formaldehyde for 10 min at room temperature before the permeabilization and the antibody incubations as described for the ovaries.

### Western blotting

10 to 20 μl of freshly dissected ovaries was collected in cold PBS and snap-frozen. The ovaries were homogenised in a 10-times volume of the RIPA buffer (50 mM Tris-HCl pH 7.5, 150 mM NaCl, 1% TritonX-100, 0.1% SDS, 0.1% Na-deoxycholate, 1 mM EDTA, and 1x cOmplete protease Inhibitors (Roche)) on ice. The lysate was cleared by centrifugation, boiled in Laemmli buffer before loading onto the SDS-PAGE. The gel was transferred onto a Nitrocellulose membrane and incubated with the primary antibodies in the following dilutions: 1:500 for anti *D. pseudoobscura* Piwi, Aubergine, and Ago3, anti *D-eugracilis* Piwi and Aubergine. The HPR-conjugated secondary antibodies were used for the standard ECL detection.

### RNA in-situ fluorescent hybridisation (FISH)

RNA FISH protocol was modified from Andersen et al^36^. Briefly, short oligo DNAs for each target (see Supplementary Table 6) were pooled and labelled with 5-Propargylamino-ddUTP-Cy3 or -Cy5 (Jena Biosciences) using the Terminal Deoxynucleotidyl Transferase (Thermo Fisher)^52^, which yielded at least 50% labelling efficiencies. Ovaries were freshly dissected from 2 - 7d old females and fixed in PBS containing 4% formaldehyde. The fixed ovaries were permeabilised overnight at 4°C in 70% ethanol and washed twice by RNA FISH wash buffer (10 % (v/w) formamide in 2x SSC). Subsequently, the ovaries were resuspended in 50 µ L Hybridization Buffer (10 % (v/w) dextran sulfate and 10 % (v/w) formamide in 2x SSC) and incubated with 1 pmol of labelled oligo probes for overnight at 37°C. The ovaries were then washed several times in RNA FISH Wash Buffer and stained by DAPI before mounting. For the double-staining of FISH probes and antibodies, we first performed the immuno-staining and followed the same procedure of the FISH from the permeabilization step. Confocal images were taken on a Zeiss LSM 800 and processed by Fiji.

### RT-qPCR analysis of *gypsy* expression

Freshly dissected ovaries from 2 – 7d old females were homogenised in Trizol to obtain the total RNA. The total RNA was further treated with RNase-free DNase I (NEB, M0303) and reverse-transcribed using the random hexamer and Superscript II (Invitrogen/Thermo Fisher) following the manufacturer’s protocol. qPCR was performed using GoTaq DNA polymerase (Promega), EvaGreen (Biotium) and the following primers: *rp49*-fw: CCGCTTCAAGGGACAGTATCTG, *rp49*-rv: ATCTCGCCGCAGTAAACGC, *gypsy*_spliced-fw: CAACAATCTGAACCCACCAATCT, *gypsy*_spliced-rv: TATGAACATCATGAGGGTGAACG. The level of *gypsy* expression was normalised to *rp49*. The expression of *gypsy* in the wild type ovaries was below detection, lower than a second control ovaries that weakly overexpressed *gypsy* (data not shown). The level of *gypsy* expression in the Yb-depleted ovaries was thus compared to the second control in Figure 1B.

### RNA sequencing

Polyadenylated RNA was purified from the DNase-treated total RNA using the oligo d(T)25 magnetic beads (NEB, S1419) and used for the library preparation. Libraries were cloned using the NEBNext Ultra Directional II RNA Library Prep Kit for Illumina (NEB, E7760), following the manufacturer’s instruction, and amplified by the KAPA LongRange DNA polymerase (Sigma, KK3502) using the universal forward primer, Solexa_PCR-fw: (5’-AATGATACGGCGACCACCGAGATCTACACTCTTTCCCTACACGACGCTCTTCCGATCT) and the barcode-containing reverse primer TruSeq_IDX: (5’-CAAGCAGAAGACGGCATACGAG<u>A</u>Tx<u>xxxxx</u>GTGACTGGAGTTCAGACGTGTGCTCTTCCGATCT where xxxxxx is the reverse-complemented barcode sequence). QuantSeq 3’ mRNA-Seq Library Prep Kit FWD for Illumina from Lexogen was used to generate the mRNA 3’-end sequencing library from the total RNA of *Drosophila eugracilis*. Amplified libraries were multiplexed and sequenced on a HiSeq platform in the paired-end 150bp mode by GENEWIZ/Azenta.

### H2RNA sequencing analysis

Both R1 and R2 reads from the poly-A+ RNA sequencing reads, and the R1 reads from the mRNA 3’-end sequencing were trimmed of the Illumina-adapter sequences using the FASTX-Toolkit from the Hannon Lab. An additional stretch of poly-A sequences was also removed from the mRNA 3’-end sequencing reads. The trimmed reads were subsequently filtered by the sequencing quality. Only the paired and unfiltered reads were then mapped to the Nanopore assembly (Miller et al. ^51^(PMID: 30087105)) of the *D. pseuduoobscura* genome, the UCBerk_Dbif_1.0 assembly (GenBank accession: GCA_009664405.1) of the *D. bifasciata* genome, and the ASM1815383v1assembly (RefSeq accession: GCF_018153835.1) of the *D. eugracilis* genome using STAR 2.7.3a allowing up to three mismatches. The coverage of uniquely mapped reads was counted using bedtools 2.28.0 and normalised to one million genome-unique mappers.

### Chromatin immuno-precipitation (ChIP) sequencing

ChIP was performed as previously described^36^. Briefly, ∼100 µ l of ovaries were dissected into ice-cold PBS and cross-linked PBS containing 2% para-formaldehyde for 10 minutes at room temperature while mixing several times. The cross-linking was quenched by Glycine. The cross-linked ovaries were washed in PBS twice and sonicated in 1.2ml of the ChIP lysis buffer (150mM NaCl, 20mM Tric-Cl, pH 8.0, 2mM EDTA, 1% Triton-X and 0.1% SDS) using the BioRuptor (ten cycles of 30sec/30sec ON/OFF). The lysate was cleared and incubated overnight at 4°C with 1 µ g of mouse monoclonal anti RNA polymerase II CTD (gift from Hiroshi Kimura^53^). Antibodies were captured by 20 µ l of a 50:50 mix of Protein-A and Protein-G Dynabeads (Life Technologies). The beads were washed several times sequentially in the buffers containing low salt (150 mM NaCl), high salt (500 mM NaCl), 250 mM LiCl, and the TE buffer before eluting in the Elution buffer (1% SDS, 0.1 M NaHCO3, and 10 mM DTT). The eluate was treated with RNase Cocktail (Thermo Fisher, AM2286), de-crosslinked for 5 hours at 65°C. De-crosslinked samples were treated with Proteinase K and the DNA was extracted by acid-phenol: chloroform and precipitated by 2-propanol. The DNA was then End-repaired and ligated to the NEBNext adaptor for Illumina (NEB, E7337A). The libraries were amplified by KAPA polymerase using the same primers as for the RNA sequencing.

### ChIP sequencing analysis

ChIP sequencing reads were trimmed of the Illumina-adapter sequences and filtered by quality using the FASTX-Toolkit. Read 1 reads were then mapped to the *Drosophila pseudoobscura* Nanopore assembly and the UCBerk_Dbif_1.0 assembly using bowtie 1.2.3 allowing up to three mismatches. The coverage of uniquely mapped reads was counted using bedtools 2.28.0 and normalised to one million genome-unique mappers.

### Small RNA cloning

We generated small RNA libraries from 5 μg of oxidised or unoxidized total RNA using a modified protocol of the original method^54^. To prepare oxidised small RNA libraries, the size of 19 to 35nt of RNA was first selected from the total RNA by PAGE using radio-labelled 19mer spike (5’-CGUACGCGGGUUUAAACGA) and 35mer spike (5’-CUCAUCUUGGUCGUACGCGGAAUAGUUUAAACUGU). The size-selected RNA was precipitated, oxidised by sodium periodate^55^, and size-selected for the second time by PAGE. To prepare unoxidized small RNA libraries, 2S ribosomal RNA was first removed from the total RNA using a biotinylated oligo DNA from IDT (5Biotin-TEG/TACAACCCTCAACCATATGTAGTCCAAGCA), followed by the size-selection using 19mer and 35mer spikes. The size-selected small RNAs were ligated to the 3’ adapter from IDT (5rApp/NNNNAGATCGGAAGAGCACACGTCT/3ddC where Ns are randomised) using the truncated T4 RNA Ligase 2, K227Q (NEB), followed by a third and second PAGE for oxidised and unoxidized libraries, respectively. Subsequently, the RNA was ligated to the 5’ adaptor from IDT (ACACUCUUUCCCUACACGACGCUCUUCCGAUCUNNNN where Ns are randomised) using the T4 RNA Ligase 1 (NEB). Adaptor-ligated RNA was reverse-transcribed by SuperScript II and amplified by KAPA polymerase using the same primers as for the RNA sequencing.

### Preparation of embryonic small RNA libraries

Embryonic small RNA libraries were prepared from the total RNA collected from 0- to 2-hour old eggs laid by females that were reared at room temperature in a cage with apple agar plates. Embryos were bleached and immediately transferred to TriZol before preparing the total RNA, which was subsequently treated with the DNase. The size-selected RNA was oxidised before ligating the adaptors.

### Immuno-precipitation of *Drosophila eugracilis* Piwi and Aubergine for small RNA cloning

100 μl of *Drosophila eugracilis* ovaries were dissected into PBS on ice. The tissue was homogenised on ice twice in 0.3 ml of the RIPA buffer. The lysate was cleared by centrifugation and diluted with 2.4 ml of IP dilution buffer (50 mM Tris-HCl pH 7.5, 150 mM NaCl), and split into two reactions. 5 μg of antibodies against Piwi and Aubergine were each coupled to 50:50 of protein-A and protein-G Dynabeads (Life Technologies). Lysates were incubated with bead-coupled antibodies, rotating at 4 °C overnight. Subsequently, the beads were captured and washed five times with IP wash buffer (50 mM Tris-HCl pH 7.5, 500 mM NaCl, 2 mM MgCl2, 10% glycerol, 1% Empigen). For Aub IP, 150 mM NaCl was used instead of 500 mM NaCl. The IP was eluted in 10mM DTT and 0.1% SDS in 1x TE at 85 degrees. The bound RNA was extracted using acid-phenol:chloroform followed by 2-propanol precipitation, mixed with the radio-labelled spikes before the size-selection. The size-selected RNA was oxidised before ligating the adaptors.

### Analysis of *Drosophila* small RNA sequencing libraries

The R1 sequencing reads were trimmed of the Illumina-adapter sequence using the FASTX-Toolkit. The four random nucleotides at both ends of the read were further removed. The trimmed reads of 18 to 40nt in size were first mapped to the infra-structural RNAs, including ribosomal RNAs, small nucleolar RNAs, small nuclear RNAs, microRNAs, and transfer RNAs (tRNAs) using bowtie 1.2.3 allowing up to one mismatch. Sequences annotated in the dm6 r6.31 assembly of the *D. melanogaster* genome were used. microRNA reads were used for a normalisation purpose. Reads that originate from the 19mer and 35mer spikes were also removed. The remaining reads were used for all the downstream analyses. The trimmed and unfiltered reads were then mapped to the Nanopore assembly of the *D. pseuduoobscura* genome, the UCBerk_Dbif_1.0 assembly of the *D. bifasciata* genome, and the ASM1815383v1assembly of the *D. eugracilis* genome, the ASM1815372v1 assembly (RefSeq accession: GCF_018153725.1) of the *D. mojavensis* genome, and the dm6 r6.31 of the *D. melanogaster* genome using bowtie allowing up to one mismatch. Reads that mapped to the 100nt upstream and downstream genomic regions of tRNA insertions were also removed. The method of identifying tRNA gene loci in the unannotated genome is further described in the code available in the git repository. Bedtools was used to count the coverage of the mapped reads. The genome-unique mappers from the oxidised total small RNA libraries were used to visualise piRNAs expressed from the piRNA clusters.

The small RNA library (SRR1746887) of the *D. melanogaster* ovarian somatic cells (OSCs) from the previous study^13^ was analysed in the same manner as other libraries generated in this study.

For the mapping of transposon-derived small RNAs, we first selected reads that mapped at least once to the respective genomic sequences. Genome-mapped unfiltered reads were subsequently mapped to the collection of autonomous *Drosophila* transposon sequences retrieved from the RepBase in March 2022, or to the curated set of transposon sequences available for *D. melanogaster*^18^, using bowtie allowing up to three mismatches with the “--all --best --strata” option. We selected transposons in the *D. pseudoobscura* and *D. eugracilis* genomes for measuring the strand bias of piRNAs and the linkage analyses using the following criteria. We started with 150 RepBase entries that expressed most abundant piRNAs in the whole ovary samples of each species. Followingly, entries of similar sequences were removed by counting the piRNA reads that were mapped to multiple entries. This process resulted in 109 and 94 “non-redundant” transposon entries from *D. pseudoobscura* and *D. eugracilis*, respectively. For *D. melanogaster*, we started with 98 curated transposons that produced most abundant ovarian piRNAs in the *w^1118^* strain. Additionally, *mariner* transposons in *D. eugracilis* were excluded from the analysis of the piRNA strand bias because of the terminal inverted repeats. All the transposon sequences used in this study are available in the git repository.

We used the replicate 1 of the oxidised whole ovary small RNA library from *D. eugracilis* for all the analyses and additionally used the replicate 2 for the *in-trans* ping-pong analysis.

### Tile coverage analysis of the small RNA sequencing reads

We carried out two genomic tile analyses to measure the abundance of piRNAs. We only considered reads that are 23nt or longer as piRNAs. We used the genome-unique mappers and uniquely-mappable 0.5kb tiles to generate the scatter plots, and all genome mappers and 0.2kb tiles to quantify the soma-enriched piRNAs. We first identified 0.5kb tiles that are covered at least 85% by unique regions by mapping artificially made 25mers against the whole genome. We then counted the number of reads that mapped to individual uniquely-mappable 0.5kb tiles. Counts were normalised to the number of genome-unique piRNA mappers. To count the abundance of all genome piRNA mappers in the 0.2kb tiles, we first mapped the sequencing reads to the genome using bowtie with the “--all --best --strata” option and divided individual mapping instances by the number of mapping events per read, in order to evenly distribute multi-mappers across all repeats in the genome. We then counted the number of reads that mapped to individual 0.2kb tiles. “Somatic” 0.2kb tiles were defined as those tiles that expressed more than ten times piRNAs in the whole ovaries than in the embryos. Of the somatic tiles, we defined “*gypsy*” sense and antisense tiles as those tiles that are covered at least half by the *gypsy* insertions predicted by the RepeatMasker. We applied the same rule to annotate the “mRNA exon” tiles and the piRNA cluster tiles. We used the RefSeq annotations for *D. pseudoobscura*, *D. eugracilis*, and *D. mojavensis*, and the FlyBase dm6 r6.31 annotations for *D. melanogaster*. We used the RNA sequencing read coverage of greater than 1 read per kilo base per million reads (RPKM) to define mRNA exons in the *D. bifasciata* genome. The coordinates of the piRNA clusters used in this study can be found in the git repository. When multiple annotations are found in the same tile, we chose the annotations in the order of “piRNA cluster”, “*gypsy* antisense/sense”, and “mRNA exons”.

### Linkage analysis of ping-pong and phasing piRNA biogenesis

We carried out the linkage analysis of transposon piRNAs as previously described^56^. Briefly, 5’ and 3’ ends of piRNA mappers (23mer or longer) were counted at each transposon coordinates. Frequencies of the co-occurrence at the linkage positions were measured across the transposon length and weighed by the abundance of piRNAs. The linkage positions of +10 between 5’ ends of sense and antisense piRNAs, and +1 between the 3’ ends and the 5’ ends of antisense piRNAs were measured for the ping-pong and the phasing biogenesis, respectively. Frequencies were measured in the window of 20nt and the Z scores are calculated as the deviation of the frequency value at the linkage position from the mean frequency divided by the standard deviation of the frequencies. Transposons that expressed at least 10% from both strands were included, and those that expressed more than three times in the whole ovaries compared to the embryos were excluded from the ping-pong linkage analysis.

### Analysis of mouse small RNA sequencing libraries

The small RNA library of mouse pachytene stage spermatocytes from the previous study (SRR1104823) was analysed^57^ in the same way as *Drosophila* small RNA libraries. Briefly, sequencing reads were trimmed of the adaptors. Reads mapping to the infra-structural RNAs were filtered. Sequences of the mouse infra-structural RNAs were retrieved from the Ensembl GRCm39 assembly and the RepBase. Followingly, 18 to 40nt long reads were mapped to the GRCm39 genome using bowtie allowing up to one mismatch. Reads that are 23nt in size or longer were considered as piRNAs and used for the *in-trans* ping-pong analysis.

### Linkage analysis of *in-trans* ping-pong

Individual genome-mapping piRNA reads were trimmed to the following three regions: the g1 to g9 position (g1g9) and the most 3’ 9 nucleotides (last9) of the sense piRNA read; and the reverse-complemented sequence from the g2 to g10 position (g2g10_revComp) where the g1 is the 5’ end of a piRNA and the g2 is the penultimate 5’ end, and so on. The frequencies of every different 9mers from each category were counted using jellyfish 2.3.0 and normalised to one million genome-mappers. The sum of the frequencies was arbitrarily set to 1000. Products of the frequencies from the g1g9 and the g2g10_revComp were calculated for each 9mer and summed up to yield the *in-trans* ping-pong linkage value. The linkage value increases when two piRNAs form an *in-trans* ping-pong pair and the same sequence occurs at higher frequencies both in the g1g9 and the g2g10_revComp. We do not expect to observe any linkage between the last9 and the g2g10_revComp, therefore, the products between them were considered a genomic background. To estimate the proportion of the *in-trans* ping-pong pairs in the piRNA pool, we performed a simulation where we artificially added fixed number of pairs into randomly selected pool of 9mers. The artificial ping-pong pairs were set to have 100 counts per million reads (CPM). piRNA reads that are equal to or more abundant than 100 CPM make up about 10% of the whole *Drosophila* ovarian piRNA population. The simulation after including 0%, 1%, 3%, 5% and 10% of artificial ping-pong pairs showed that the *in-trans* ping-pong linkage value increases by 1 as the proportion of the ping-pong pairs increases by 1% (Figure S9B). Based on this result, we estimated the percentage of *in-trans* ping-pong pairs in biological samples by the linkage value of g1g9 * g2g10_revComp subtracted by the value of last9 * g2g10_revComp.

## Acknowledgements

We thank the Brennecke lab for housing several non-*melanogaster Drosophila* strains. We thank Kazu Mochizuki and Julius Brennecke for critical comments on the manuscript. The work was funded by the ANU start-up (to R.H.), Australian Research Councils (DP210102385 to R.H.) and the ANU Vice-Chancellor travel fellowship (to S.C).

## Data availability

Sequencing data and processed files have been deposited to Gene Expression Omnibus (GSE213383). Codes for the computational analyses are available at https://github.com/RippeiHayashi/noYb_Dspp.git.

## Author Contributions

R.H. conceived the study. R.H and S.C. designed, conducted and analysed the experiments. R.H and S.C. wrote the manuscript.

## Figure legends

**Figure S1.**
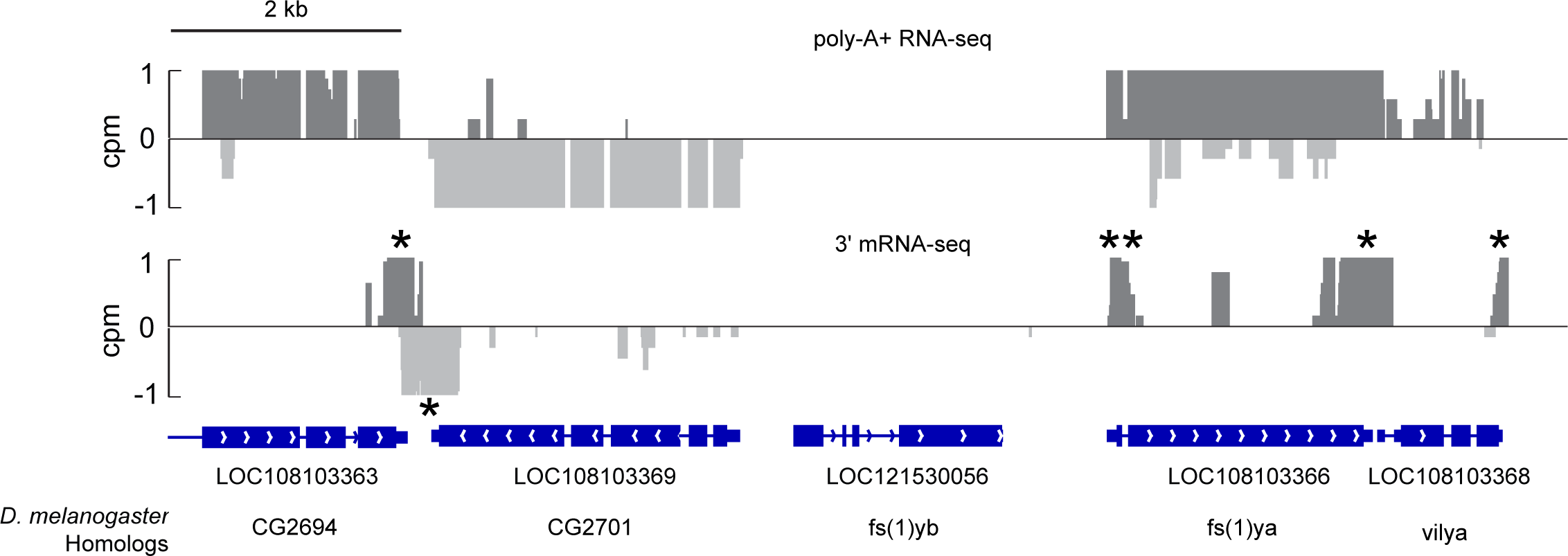
The *yb* pseudogene in *D. eugracilis* is not expressed in the ovary. Both poly-A+ RNA-seq and 3’ mRNA-seq show no expression of the pseudogene of *yb* in *D. eugracilis*. The coverage of neighbouring genes is shown in counts per million reads (CPM). Single and double asterisks indicate peaks of the 3’ mRNA-seq that correspond to canonical mRNA 3’ ends and a suspected internal priming, respectively.

**Figure S2.**
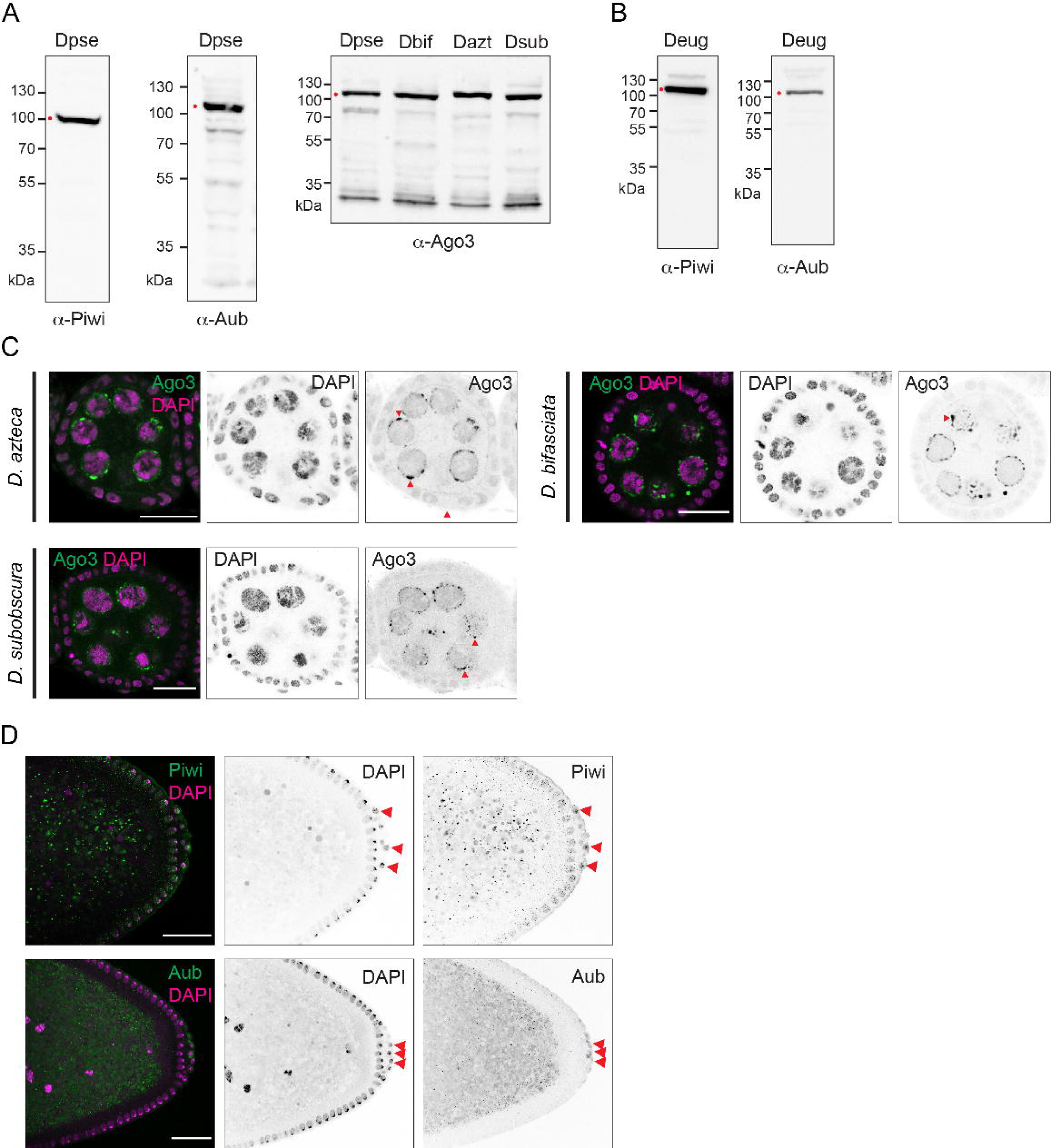
PIWI protein localisation in the egg chambers of the *obscura* group ovaries and the blastderm stage of the *D. eugracilis* embryos. **(A) and (B)** Antibodies raised against peptides from *D. pseudoobscura* and *D. eugracilis* PIWI proteins were used in (A) and (B), respectively. The lysates were prepared from ovaries from indicated species and blotted for antibodies against Piwi, Aubergine (Aub) and Ago3. Dominant bands, as indicated by red dots, were detected at predicted sizes (100 to 110 kDa) in all cases. Species names are abbreviated as follows: *D. pseudoobscura*; Dpse, *D. bifasciata*; Dbif, *D. azteca*; Dazt, *D. subobscura*; Dsub. and *D. eugracilis*; Deug. **(C)** Immuno-fluorescent stainings of (Ago3 in green and DAPI in magenta) of *D. azteca*, *D. bifasciata*, and *D. subobscura* egg chambers show a peri-nuclear localisation of Ago3 in all three species. Peri-nuclear granules, as indicated by arrowheads, are seen in all species. **(D)** Immuno-fluorescent staining of Piwi and Aubergine in *D. eugracilis* mid-blastoderm stage embryos showing the localisation of Piwi and Aubergine in the pole cells (marked by arrowheads) and in the zygotic somatic nuclei (Piwi only). Embryos of 1 – 1.5hr post fertilisation were stained. This is the stage after the nuclei reach the periphery of the blastoderm before the nuclear elongation and the onset of cellularisation. The bulk of zygotic transcription has not started in this stage. Scale bars = 20 μm

**Figure S3.**
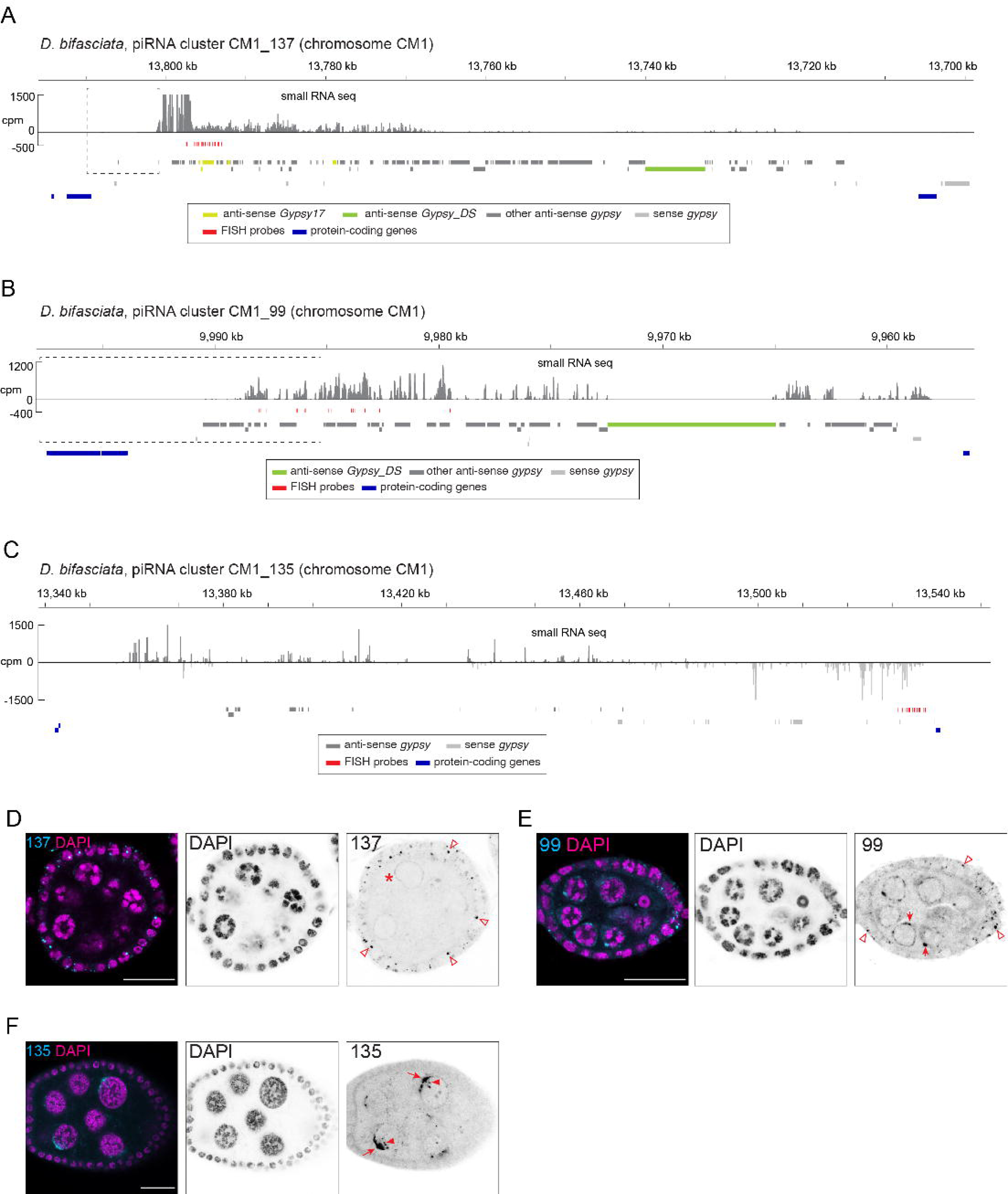
piRNA clusters in *D. bifasciata* ovaries. **(A) - (C)** Shown are the coverage of piRNA reads (>22nt) in counts per million genome mappers (CPM) from the oxidised whole ovary small RNA library of *D. bifasciata* that uniquely mapped to the cluster regions. Sense and antisense reads are coloured in dark and light gray, respectively. Coloured bars indicate *gypsy* insertions predicted by RepeatMasker, annotated protein-coding mRNA exons and the FISH probes. Dotted box in (A) and (B) indicate the putative transcription start sites of the cluster, for which magnified views are shown in Figures S4B and C. **(D) - (F)** RNA FISH against transcripts from the piRNA clusters. Somatic signals as indicated by open arrows are seen for the FISH against clusters CM1_137 (D) and CM1_99 (E), while the FISH against the cluster CM1_135 (F) only stains the germline cells. The FISH against CM1_137 weakly stained the germline at the nuclear periphery (asterisk) while distinct peri-nuclear puncta are seen in the FISH against CM1_99 and CM1_135 (arrows). Putative sites of transcription in the nuclei as indicated by arrowheads are only seen in CM1_135.

**Figure S4.**
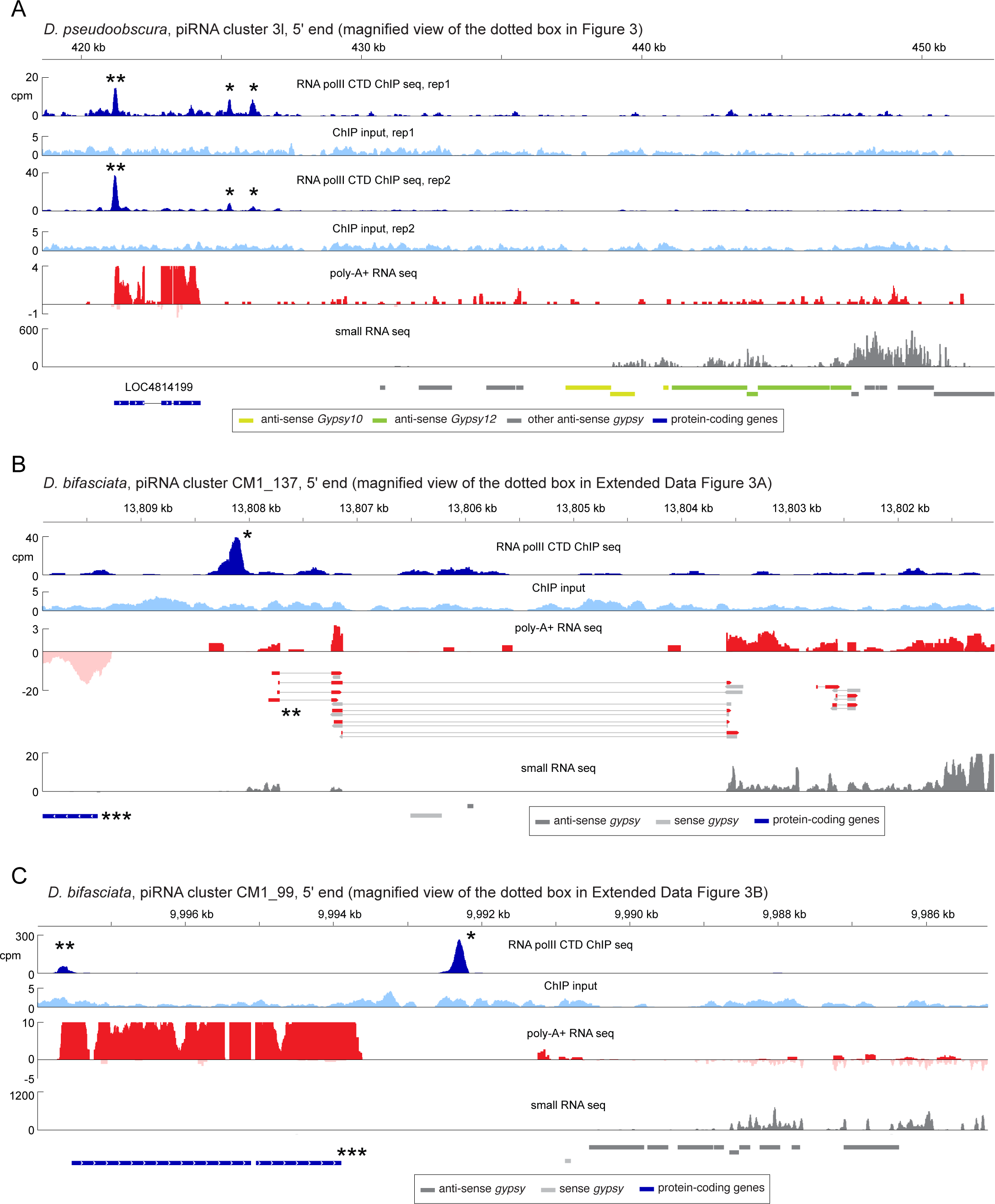
Somatic piRNA clusters in the *obscura* group resemble protein-coding genes. **(A) - (C)** Shown are the coverage of RNA polymerase II ChIP-seq (blue), and its input controls (light blue), poly-A+ RNA-seq (sense in red and antisense in pink), and the piRNA reads (>22nt) at the 5’ end of the somatic piRNA clusters. Y-axes indicate counts per million genome mappers (CPM) for all tracks. RNA-seq read pairs that span introns are indicated as red and gray bars in pairs. Coloured bars at the bottom indicate *gypsy* insertions and annotated protein-coding mRNA exons. Peaks of RNA polymerase II in front of the clusters and at the neighbouring gene promoter regions are marked by single and double asterisks, respectively. Triple asterisks indicate protein-coding exons in *D. bifasciata* predicted by homology to *D. pseudoobscura* proteins.

**Figure S5.**
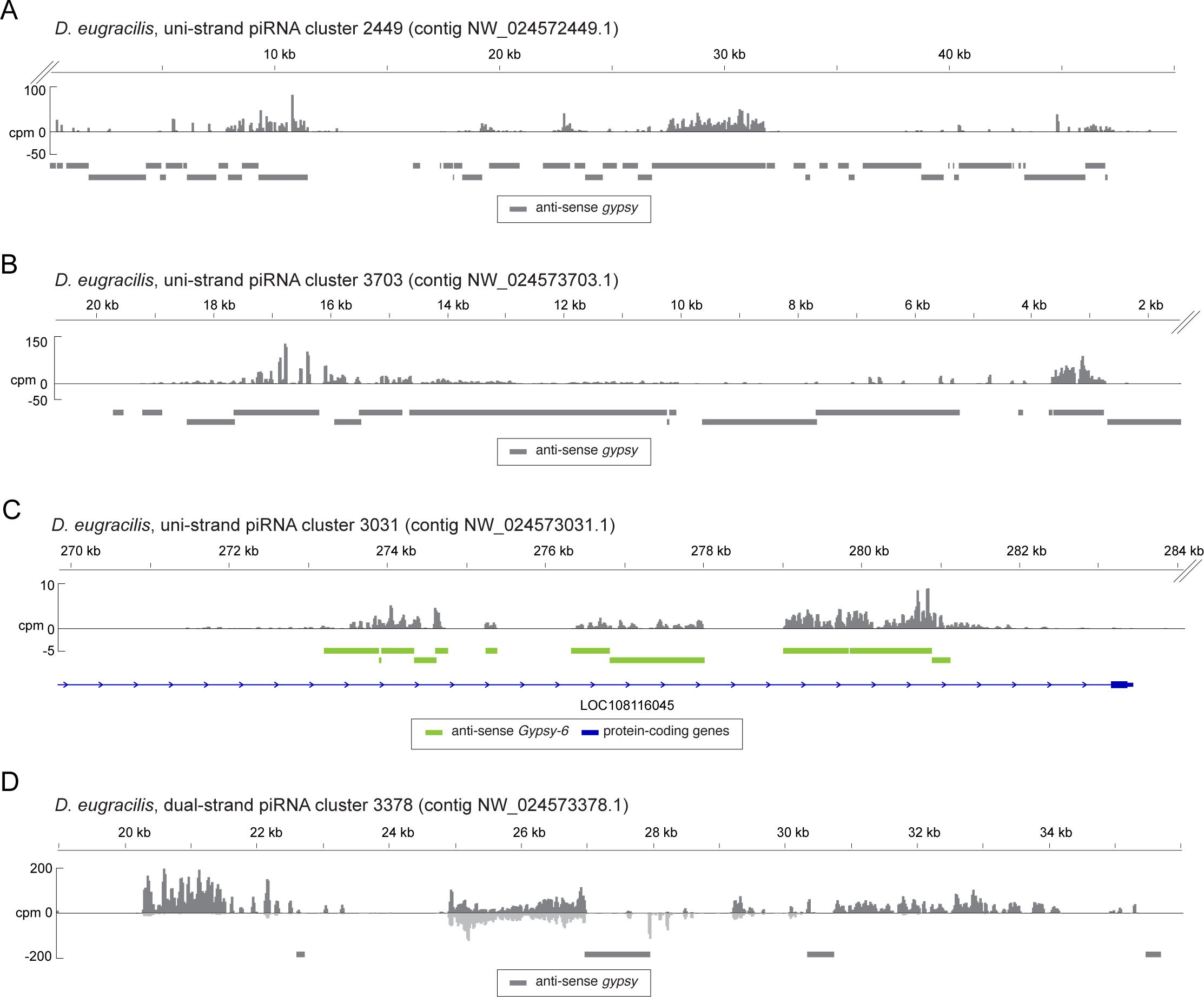
piRNA clusters in the *D. eugracilis* genome. **(A) - (D)** Shown are the coverage of piRNA reads (>22nt) in counts per million genome mappers (CPM) from the oxidised whole ovary small RNA library of *D. eugracilis* that uniquely mapped to the cluster regions. Sense and antisense reads are coloured in dark and light gray, respectively. Coloured bars indicate *gypsy* insertions predicted by RepeatMasker, and an annotated protein-coding mRNA exon. The entirety of the uni-stranded clusters could not be determined because they are found at the end (indicated by double dashed lines) of the chromosome contigs.

**Figure S6.**
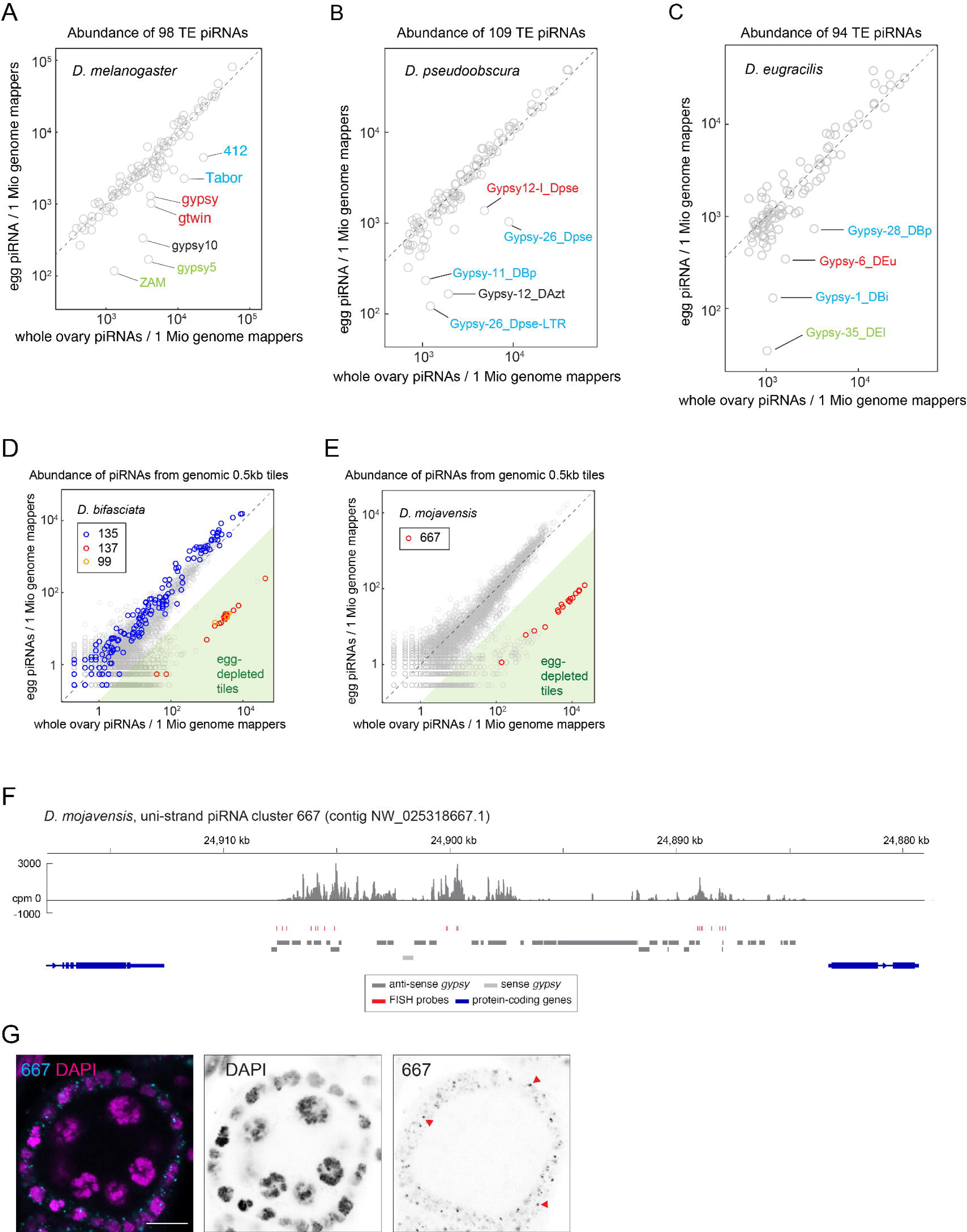
Characterisation of somatic piRNA expression in Drosophila species. **(A) - (C)** Abundance of ovarian (X axes) and embryonic (Y axes) piRNAs mapping to individual transposons in *D. melanogaster* (A), *D. pseudoobscura* (B) and *D. eugracilis* (C) genomes. Transposons that expressed piRNAs more than three times in the ovaries than in the embryos are marked. Colours indicate families within the “errantiviridae/412” group of Ty3/Gypsy superfamily: red; group “Gypsy”, green; group “17.6”, cyan; group “412/mdg1”, and black; unclassified. **(D and E)** Scatter plots showing the abundance of piRNAs from the whole ovaries (X axis) and the eggs (Y axis) that uniquely mapped to the individual 0.5 kb tiles of the *D. bifasciata* genome in (D) and *D. mojavensis* genome in (E). The dual-stranded germline clusters are coloured in blue while uni-stranded somatic clusters are coloured in orange and red. Tiles that expressed piRNAs more than ten times in the whole ovaries than in the embryos are shaded in green. **(F)** Shown is the coverage of piRNA reads (>22nt) in counts per million genome mappers (CPM) from the oxidised whole ovary small RNA library of *D. mojavensis* that uniquely mapped to the cluster region. Sense and antisense reads are coloured in dark and light gray, respectively. Coloured bars indicate sense and antisense *gypsy* insertions predicted by RepeatMasker, annotated protein-coding mRNA exons, and FISH probes. (**G)** RNA FISH, showing the expression of the cluster 667 in the somatic cells of a *D. mojavensis* egg chamber. Focused signals in the somatic cells are indicated by arrowheads. Scale bars = 10 μm

**Figure S7.**
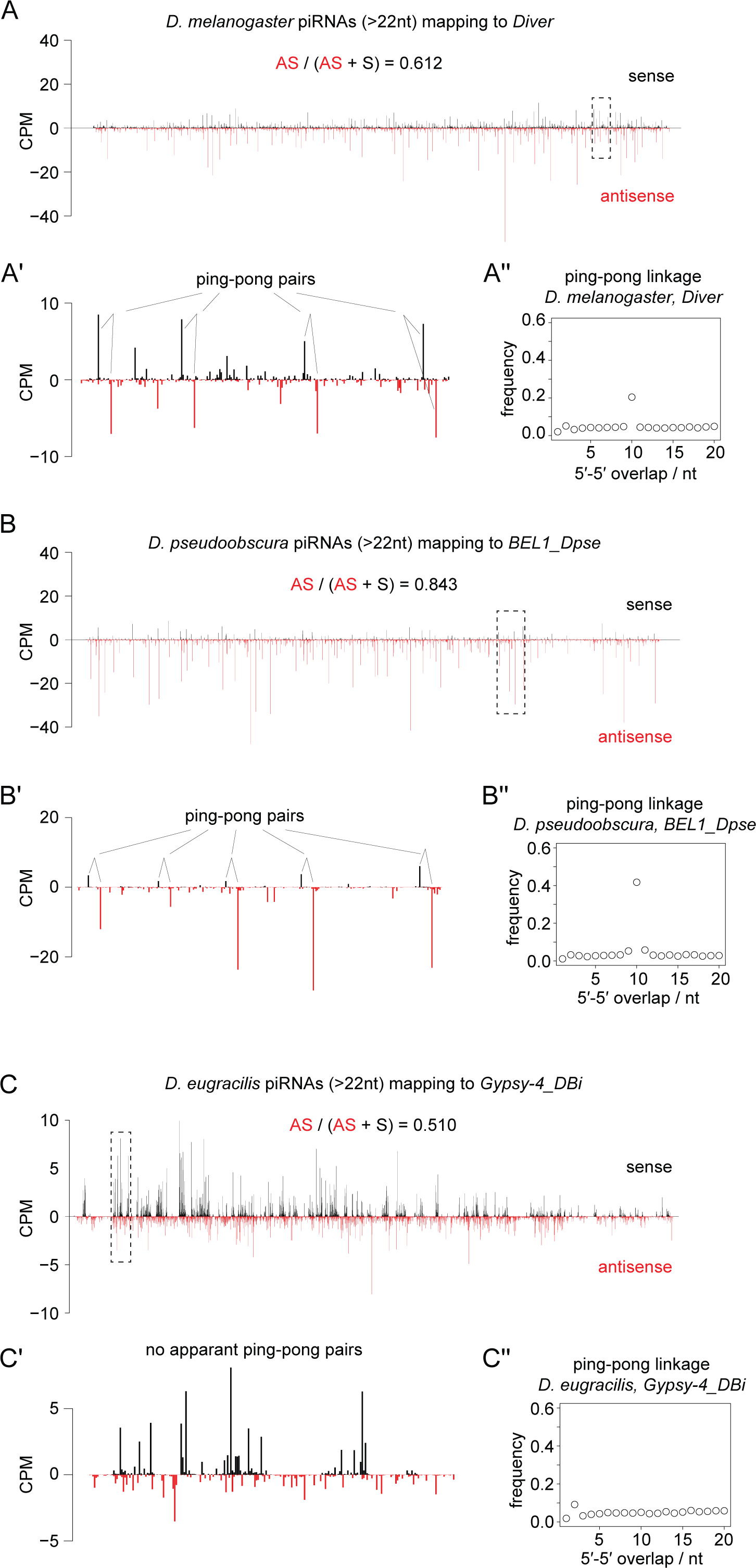
Absence of ping-pong transposon piRNAs in *D. eugracilis* ovaries. **(A) – (C)** Shown are the 5’ end coverage of piRNA reads (>22nt) from *D. melanogaster* (top), *D. pseudoobscura* (middle) and *D. eugracilis* (bottom) ovaries mapping to indicated transposon sequences in counts per million genome mappers (CPM). Sense and antisense reads are coloured in black and red, respectively. Putative ping-pong pairs or absence of them are highlighted in the magnified view of the regions shown by dashed boxes (A’, B’, and C’). Frequencies of the 5’ overlapping bases between sense and antisense piRNAs are calculated where the characteristic 10nt overlap is visible when there is a prominent ping-pong (A’’, B’’, and C’’).

**Figure S8.**
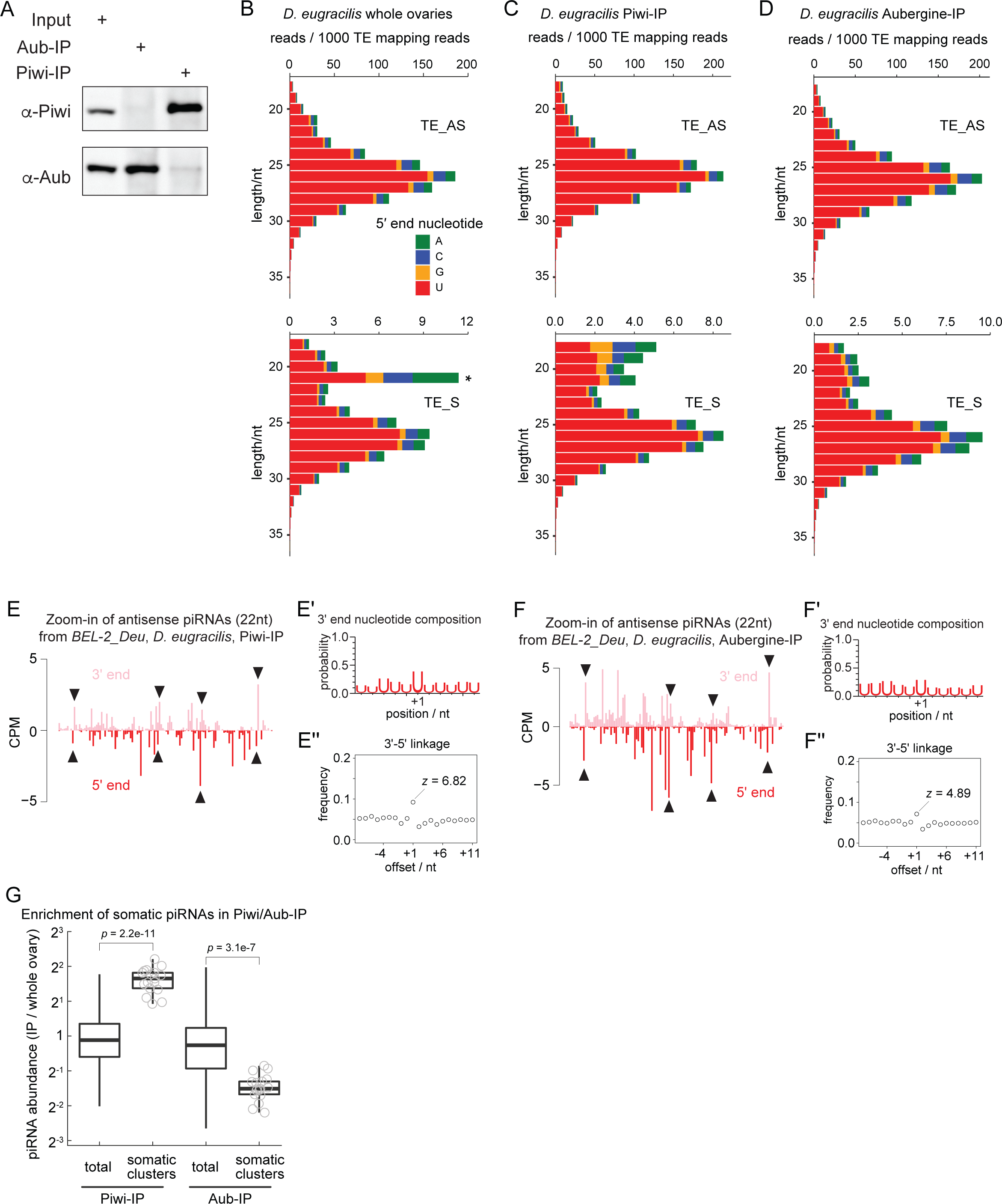
*D. eugracilis* Piwi and Aubergine are bound to nearly identical pools of piRNAs. **(A)** A western blotting showing the specificities of antibodies against *D. eugracilis* Piwi and Aubergine (Aub). Input ovary lysates and the elutions after the immuno-precipitation (IP) are loaded and blotted against respective antibodies. **(B – D)** Size distribution of total ovarian (B), Piwi- (C) and Aubergine-bound (D) small RNAs mapping to transposon sense (bottom) and antisense (top) sequences. The proportion of nucleotides at the 5’ end is shown by different colours, showing the Uridine preference. The abundance is normalised to the total transposon mapping reads, showing that the majority of reads are antisense for all three libraries. There are very few putative siRNAs (21nt, marked by asterisk) compared to piRNAs (24 to 29nt), that are only detected in the sense reads and depleted in the IP libraries. **(E and F)** Shown are the 5’ and 3’ ends of Piwi- (E) and Aubergine-bound (F) piRNAs mapping to the antisense strand of *BEL-2_Deu* from the region indicated by a dashed box in Figure 6A. The 3’ and 5’ ends of piRNAs that are one nucleotide apart, hence the putative products of phasing, are marked by arrowheads. **(E’ and F’)** Shown are the frequencies of Uridines found at positions relative to the 3’ ends of piRNAs mapping to *BEL-2_Deu*. +1 corresponds to the immediate downstream nucleotide position. **(E’’ and F’’)** Shown are the frequency plot of the 3’-5’ linkage of antisense *BEL-2_Deu* piRNAs. The z scores of the linkage position +1 are shown. **(G)** A box plot showing the relative abundance of genome-unique piRNAs from *D. eugracilis* mapping to 0.5kb tiles, comparing the whole ovaries and Piwi- and Aubergine-bound pools. piRNAs mapping to the tiles from somatic clusters are enriched and depleted in Piwi- and Aubergine-bound pools, respectively. *p*-values are calculated by Mann-Whitney U test.

**Figure S9.**
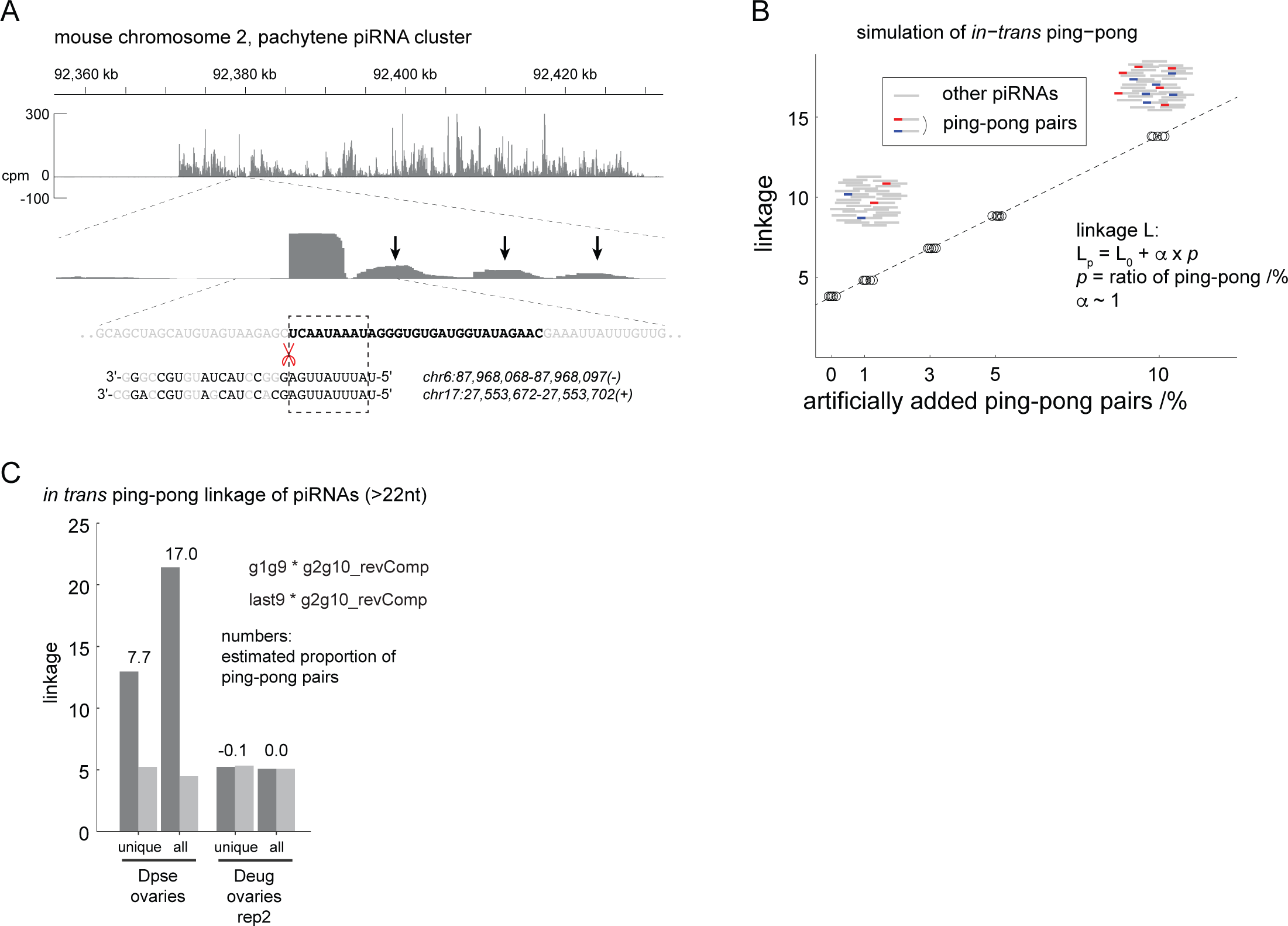
*in-trans* ping-pong of simulated, mouse pachytene, and Drosophila piRNAs. **(A)** Shown at the top is the coverage of piRNA reads (>22nt) from a mouse pachytene piRNA cluster at chromosome 2 (PMID: 26115953). Shown at the bottom is an example of putative phasing events. The most abundant piRNA in this region is likely made as a result of slicing by piRNAs from distant genomic loci, which is followed by production of downstream piRNAs as indicated by arrows. **(B)** Shown are the *in-trans* ping-pong linkage values of simulated random piRNA pools with varying extent of ping-pong pairs artificially included (five replicates each). The linkage value increases by one as the proportion of artificially-added ping-pong pairs increases by one percent. **(C)** Shown are *in-trans* ping-pong linkage values of genome unique piRNAs and all piRNA mappers from *D. pseudoobscura* and the second replicate of *D. eugracilis* ovarian small RNA libraries. Estimated proportions of *in-trans* ping-pong pairs out of all piRNAs are shown in percentage.

**Figure S10.**
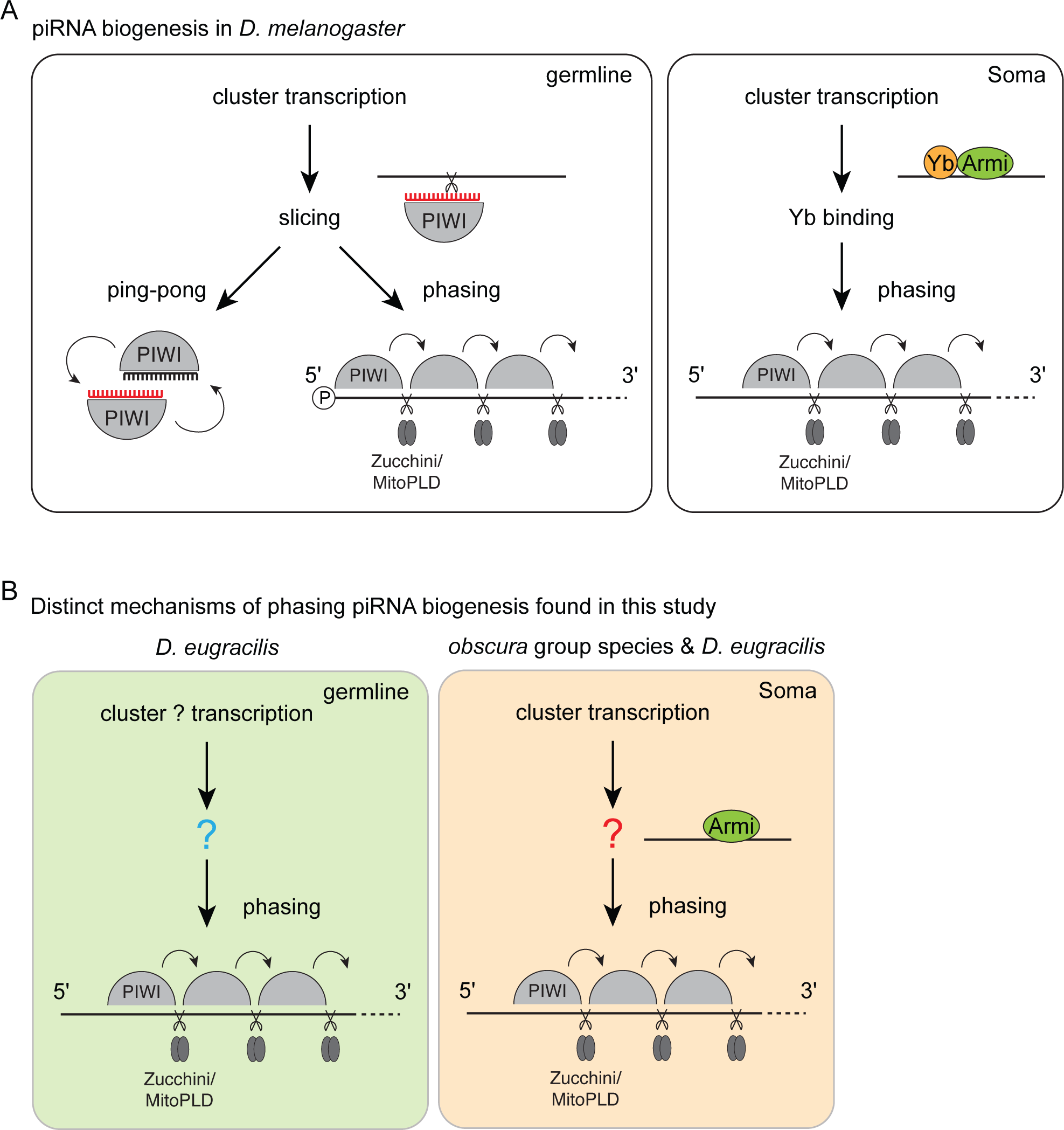
piRNAs biogenesis mechanisms in *Drosophila*. **(A)** piRNAs are made by ping-pong and phasing where ping-pong is specific to the germline and phasing can be found in both germline and soma compartments of *D. melanogaster* ovaries. Phasing is initiated by a slicing event in the germline, which also triggers the ping-pong loop, whereas Yb is required for recruiting Armitage (Armi) to the cluster-derived transcripts for phasing in the soma. **(B)** Shown are novel mechanisms of phasing piRNA biogenesis found in this study. piRNAs are predominantly made by phasing without slicing events in the *D. eugracilis* germline while Yb is dispensable for an efficient processing of cluster-derived transcripts in the somatic tissue of *obscura* group species and *D. eugracilis*.

**Figure S11.**
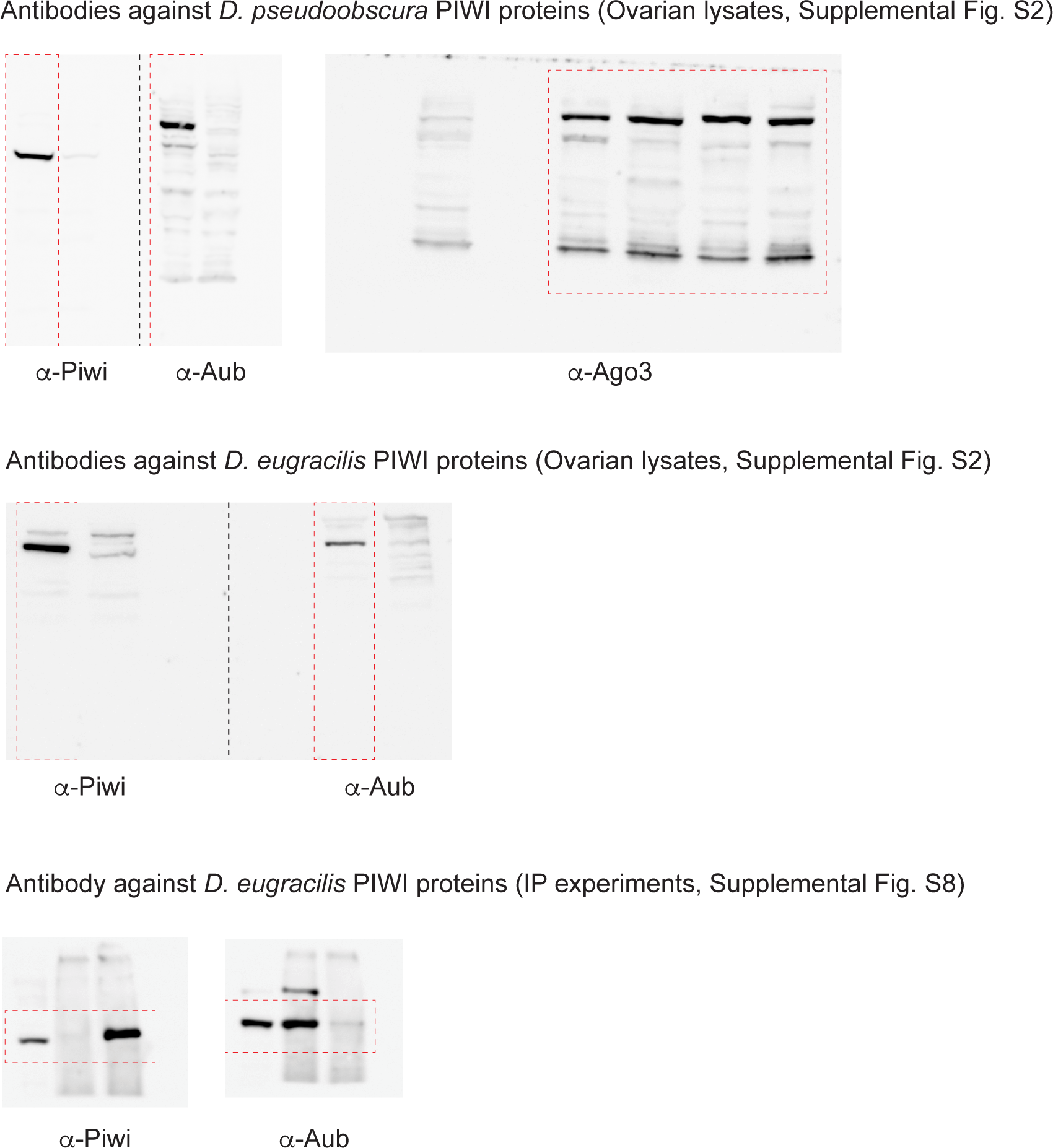
Raw images of the western blotting. Regions that are used for other figures are indicated by red dashed boxes.

**Supplementary Table 1**

**The list of genome assemblies of Drosophila species examined for the conservation of *yb***

The first tab lists GenBank and RefSeq genome assemblies and the second tab lists Nanopore assemblies from Miller et al ^51^.

**Supplementary Table 2**

**Comparison of Env protein sequences from Gypsy_DM, Gypsy12_Dpse and Gypsy_DS**

(A) Amino acid sequences of Env proteins from *Gypsy_DM*, *Gypsy12_Dpse* and *Gypsy_DS* are shown. Sequences of *Gypsy12_Dpse* and *Gypsy_DS* Env are taken from the insertions found in the *D. pseudoobscura* and the *D.bifasciata* genomes, respectively. (B) and (C) Alignments of Gypsy Env protein sequences made by Clustal Omega are shown. Functionally important motifs are shown in boxes; from the N-termius, signal peptide, furin cleavage site, and the transmembrane domain ^60, 61^. *Gypsy_DS* Env protein was previously suspected to lack the peptide signal and the transmembrane domain ^60^. However, the genomic insertion shown here retains those motifs.

**Supplementary Table 3**

**The list of intact copies of *gypsy-env* retrotransposons in the *D. pseudoobscura*, *D. bifasciata*, *D. azteca*, and *D. eugracilis* genomes.**

**Supplementary Table 4**

***aubergene* homologs found in the *obscura* group species**

**Supplementary Table 5**

**Composition of somatic piRNAs across Drosophila species**

The abundance of different classes of somatic piRNAs (>22nt) was measured by intersecting the mappings to the genome annotations. Numbers are used to make the bar chart shown in Figure 6.

**Supplementary Table 6**

**Sequences of oligo DNA probes used for RNA FISH experiments**

