## Supplementary_TableS2 for "Mechanistic divergence of piRNA biogenesis in Drosophila"

```
Supplementary Table 2

A.

>GYPSY_I_3p
MMFIPLVVANARITDFSHANYIPVLDGDVLVFEQRDLLKHSSNLSEYASMIDETQKLSESFPHSHMRKLLEVDTDHLRTLLSVLKVHHRIARSLDFLGTALKVVAGTPDATDLFKIKITEAQLVESNSRQIAINSETQKQINKLTDTINKVINARKGDLVDTPHLYEALLARNRMLSTEIQNLILTITLVKSNIINPTILDHADLKPLVEQDTPIVSLIEASKIRVLQSENSIHILIAYPRVKFSCKKVAVYPVSHQHTILRLDEDTLAECEHDTFAVTGCTDTTHFTFCERSRRETCVRSLHAGNAAQCHTQPSHLREINPVDDGVVIINEAAAHVSTDGSPETLIEGTYLVTFERTATINGSEFVNLRKTLSKQPGIVRSPLLNIVGHDPVLSIPLLHRMSNENLHSIQNLMDDVESEGSPRLWFVAGVVLNFGLIGSLALYLALRRRRASREIQRTIDTFNMTEDGHKLEGG

GCF_009870125.1_UCI_Dpse_MV25 (NC_046679.1 30539870 30541304 -)
>Gypsy12_Dpse_env_30541304
MIILLVALVNARITDYSHSDYVPILDGDILVWDEINYLRHSTNLTDYERMADETANLTEMFPQSHMRKLLVVDTDHIRNMLATISVHHRVARSLNILGSVLKVVAGTPDADDLEKIRINEAQLIESNNRQISINSKSQEQINRLTDSVNKLLEAAKGKQIDSAHLYETLLARNRMLASELSNLMLTISLAKVNVINPVILDHDDLNSIFSNQLTNVIVTNILEVSKIKVFQSNSIIHFVIQFPKIKYICKKITIFPVAHNGTVLRLDDNIVADCNDQIVTVSDCKQTTTTTFCETSTKDSCAQGLYSGGVAHCQSQPSHLSAITLVDDGIIIINDHPAAVSFDGSAALNISGTHLITFNDYAVINGSRYQNRKNVQSRYPGVASSPLLNVTEHKRVLSLPFLHQLSEENLNFIKEIKEEVSSRSRPIFAFCLGLGICGLVCGMAMLRLYLTKKRDARQINGLMARLSAPGTATAQGGE

GCA_009664405.1_UCBerk_Dbif_1.0 (CM019041.1 9970475 9971896 +)
>Gypsy_DS_Dbif-env
LLVVLAAVNARITDFSHANYIPVLDGEVLVFDQRSYLRHSSNISEFISMIDETEKLSDSFPQSHMRKLLDVDTDHLRTLLSVLQVHHRFARSLDFLGTALKVVAGTPDASDFLKVRVTEAQLVESNSKQIIINSETQKQINRLTDTINKIISSRKGDLVDTPHLFETLLARNRILNTEIQNLILTITLAKANIVNPTILDHADLKSLIEQDTPIVSLLEASKIKVLQSENIIHILIAYPKVEFKCQKVSVYPVSHQQTILRLDEDTLAECERDTFAVTGCTVTTHNTFCERARRETCASSLHAGNTANCHTQPSHLNAIMPIDDGVVVINEATARVRTDDGAEVTVSGTFLITFERSAAINGTEFINLRKAPSKQPGTVRSPLLNIIGHDPALSIPLLHRMNINNLQSILDFKEEVIAAGSPKFWFAVGAVLNVGLICSFILFMALRRKRASLRIQKALDNFNMTEDGHHSEGG


B.

Gypsy_I_3p                     MMFIPLVVANARITDFSHANYIPVLDGDVLVFEQRDLLKHSSNLSEYASMIDETQKLSES	60
Gypsy12_Dpse_env_30541304      MIILLVALVNARITDYSHSDYVPILDGDILVWDEINYLRHSTNLTDYERMADETANLTEM	60
                               *::: :.:.******:**::*:*:****:**::: : *:**:**::*  * *** :*:*

Gypsy_I_3p                     FPHSHMRKLLEVDTDHLRTLLSVLKVHHRIARSLDFLGTALKVVAGTPDATDLFKIKITE	120
Gypsy12_Dpse_env_30541304      FPQSHMRKLLVVDTDHIRNMLATISVHHRVARSLNILGSVLKVVAGTPDADDLEKIRINE	120
                               **:******* *****:*.:*:.:.****:****::**:.********** ** **:*.*

Gypsy_I_3p                     AQLVESNSRQIAINSETQKQINKLTDTINKVINARKGDLVDTPHLYEALLARNRMLSTEI	180
Gypsy12_Dpse_env_30541304      AQLIESNNRQISINSKSQEQINRLTDSVNKLLEAAKGKQIDSAHLYETLLARNRMLASEL	180
                               ***:***.***:***::*:***:***::**:::* **. :*: ****:********::*:

Gypsy_I_3p                     QNLILTITLVKSNIINPTILDHADLKPLVEQDT---PIVSLIEASKIRVLQSENSIHILI	237
Gypsy12_Dpse_env_30541304      SNLMLTISLAKVNVINPVILDHDDLNSIFSNQLTNVIVTNILEVSKIKVFQSNSIIHFVI	240
                               .**:***:*.* *:***.**** **: :..::     :..::*.***:*:**:. **::*

Gypsy_I_3p                     AYPRVKFSCKKVAVYPVSHQHTILRLDEDTLAECEHDTFAVTGCTDTTHFTFCERSRRET	297
Gypsy12_Dpse_env_30541304      QFPKIKYICKKITIFPVAHNGTVLRLDDNIVADCNDQIVTVSDCKQTTTTTFCETSTKDS	300
                                :*::*: ***::::**:*: *:****:: :*:*:.: .:*:.*.:**  **** * :::

Gypsy_I_3p                     CVRSLHAGNAAQCHTQPSHLREINPVDDGVVIINEAAAHVSTDGSPETLIEGTYLVTFER	357
Gypsy12_Dpse_env_30541304      CAQGLYSGGVAHCQSQPSHLSAITLVDDGIIIINDHPAAVSFDGSAALNISGTHLITFND	360
                               *.:.*::*..*:*::*****  *. ****::***:  * ** ***    *.**:*:**:

Gypsy_I_3p                     TATINGSEFVNLRKTLSKQPGIVRSPLLNIVGHDPVLSIPLLHRMSNENLHSIQNLMDDV	417
Gypsy12_Dpse_env_30541304      YAVINGSRYQNRKNVQSRYPGVASSPLLNVTEHKRVLSLPFLHQLSEENLNFIKEIKEEV	420
                                *.****.: * ::. *: **:. *****:. *. ***:*:**::*:***: *::: ::*

Gypsy_I_3p                     ESEGSPRLWFVAGVVLNFGLIGSLALYLALRRRRASREIQRTIDTFNMTEDGHKLEGG-	475
Gypsy12_Dpse_env_30541304      SSRSRPIFAFCLGLGICGLVCGMAMLRLYLTKKRDARQINGLMARLSAPGTA-TAQGGE	478
                               .*.. * : *  *: :   : *   * * * ::* :*:*:  :  :.    . . :**

C.

GYPSY_I_3p             MMFIPLVVANARITDFSHANYIPVLDGDVLVFEQRDLLKHSSNLSEYASMIDETQKLSES	60
Gypsy_DS_Dbif-env      -LLVVLAAVNARITDFSHANYIPVLDGEVLVFDQRSYLRHSSNISEFISMIDETEKLSDS	59
                        ::: *...******************:****:**. *:****:**: ******:***:*

GYPSY_I_3p             FPHSHMRKLLEVDTDHLRTLLSVLKVHHRIARSLDFLGTALKVVAGTPDATDLFKIKITE	120
Gypsy_DS_Dbif-env      FPQSHMRKLLDVDTDHLRTLLSVLQVHHRFARSLDFLGTALKVVAGTPDASDFLKVRVTE	119
                       **:*******:*************:****:********************:*::*:::**

GYPSY_I_3p             AQLVESNSRQIAINSETQKQINKLTDTINKVINARKGDLVDTPHLYEALLARNRMLSTEI	180
Gypsy_DS_Dbif-env      AQLVESNSKQIIINSETQKQINRLTDTINKIISSRKGDLVDTPHLFETLLARNRILNTEI	179
                       ********:** **********:*******:*.:***********:*:******:*.***

GYPSY_I_3p             QNLILTITLVKSNIINPTILDHADLKPLVEQDTPIVSLIEASKIRVLQSENSIHILIAYP	240
Gypsy_DS_Dbif-env      QNLILTITLAKANIVNPTILDHADLKSLIEQDTPIVSLLEASKIKVLQSENIIHILIAYP	239
                       *********.*:**:*********** *:*********:*****:****** ********

GYPSY_I_3p             RVKFSCKKVAVYPVSHQHTILRLDEDTLAECEHDTFAVTGCTDTTHFTFCERSRRETCVR	300
Gypsy_DS_Dbif-env      KVEFKCQKVSVYPVSHQQTILRLDEDTLAECERDTFAVTGCTVTTHNTFCERARRETCAS	299
                       :*:*.*:**:*******:**************:********* *** *****:*****.

GYPSY_I_3p             SLHAGNAAQCHTQPSHLREINPVDDGVVIINEAAAHVSTDGSPETLIEGTYLVTFERTAT	360
Gypsy_DS_Dbif-env      SLHAGNTANCHTQPSHLNAIMPIDDGVVVINEATARVRTDDGAEVTVSGTFLITFERSAA	359
                       ******:*:********. * *:*****:****:*:* **.. *. :.**:*:****:*:

GYPSY_I_3p             INGSEFVNLRKTLSKQPGIVRSPLLNIVGHDPVLSIPLLHRMSNENLHSIQNLMDDVESE	420
Gypsy_DS_Dbif-env      INGTEFINLRKAPSKQPGTVRSPLLNIIGHDPALSIPLLHRMNINNLQSILDFKEEVIAA	419
                       ***:**:****: ***** ********:****.*********. :**:** :: ::* :

GYPSY_I_3p             GSPRLWFVAGVVLNFGLIGSLALYLALRRRRASREIQRTIDTFNMTEDGHKLEGG	475
Gypsy_DS_Dbif-env      GSPKFWFAVGAVLNVGLICSFILFMALRRKRASLRIQKALDNFNMTEDGHHSEGG	474
                       ***::**..*.***.*** *: *::****:*** .**:::*.********: ***
```
